# DNA methylation variability defines a fundamental dimension of tumor epigenomes linked to genomic instability, tumor aggressiveness, and clinical outcomes

**DOI:** 10.64898/2026.03.12.711303

**Authors:** Djansel Bukovec, Blagojche Gjorgjioski, Monika Simjanoska Misheva, Goran Kungulovski

**Author notes:** Corresponding author Dr. Goran Kungulovski, Fingerprint Diagnostics LLC, Ivan Agovski 7-1, 1000, Skopje, Republic of Macedonia.

## Abstract

**Background:** Tumors exhibit substantial cellular and molecular diversity driven by genetic and epigenetic mechanisms. Large-scale profiling efforts have established aberrant DNA methylation as a universal hallmark of cancer. Beyond changes in mean methylation levels, tumor tissues exhibit elevated DNA methylation variability at specific genomic regions within and across tumors. This constitutes a fundamental dimension of cancer epigenomes, reflecting disrupted maintenance of epigenomic states and stochastic drift, which may enable adaptation to the microenvironment, phenotypic plasticity, invasion, disease progression, and treatment resistance. However, the genome-wide organization and functional consequences of DNA methylation variability across cancer types remain incompletely understood.

**Methods:** We analyzed paired tumor–normal DNA methylation profiles across 16 cancer types to systematically quantify DNA methylation variability. Pan-cancer DNA methylation variability was consistently observed using complementary statistical approaches and multiple modes of data representation. We identified cancer-specific and pan-cancer differentially variable regions and evaluated their associations with genomic features, transcriptional and chromatin regulators, and biological processes. Variability was quantified using three measures per sample: the proportion of intermediately methylated sites (PIM), genome-wide Shannon entropy, and a DNA methylation–based stemness index. Associations with genomic instability, tumor biological features, and clinical outcomes were subsequently assessed.

**Results:** Tumor samples consistently exhibited higher DNA methylation variability than matched normal tissues, reflected by increased dispersion and wider interquartile ranges. Pan-cancer variably methylated regions were depleted in promoters and enriched in open sea regions, in heterochromatic H3K27me3-decorated PRC2-repressed domains, and at enhancers. They preferentially contained motifs for transcription factors involved in developmental regulation. Elevated DNA methylation variability, captured by higher PIM, entropy, and stemness scores, was associated with increased genomic instability manifested by higher aneuploidy, increased DNA break points, a greater fraction of the genome altered, and increased tumor mutational burden, as well as with aggressive tumor features such as lymph node involvement, post-therapy neoplasm events, and elevated hypoxia scores. Importantly, tumors with high DNA methylation variability exhibited significantly worse overall, progression-free, and disease-free survival.

**Conclusions:** DNA methylation variability is a pervasive and clinically relevant feature of tumor epigenomes, reflecting epigenetic and genetic instability, expanded regulatory plasticity, and tumor aggressiveness.

## BACKGROUND

Tumors exhibit substantial cellular and molecular diversity driven by genetic and non-genetic (epigenetic) mechanisms (1,2). Among these, DNA methylation is a central epigenetic regulator of gene expression, cellular identity, and genome stability, classically characterized by promoter-specific hypermethylation and global hypomethylation in cancer (3,4). Large-scale profiling efforts, particularly through The Cancer Genome Atlas (TCGA), have established aberrant DNA methylation as a universal hallmark of cancer and revealed distinct tumor subgroups defined by their methylation landscapes across tissues and lineages (5–7).

But in the past years, cancer has been increasingly recognized as a disease of epigenomic disorganization, in which epigenetic alterations, including DNA methylation, accompany and reinforce genetic, transcriptional, and phenotypic dysregulation (8–10). Beyond changes in mean methylation levels, accumulating evidence indicates that cancer tissues exhibit elevated DNA methylation variability at specific genomic regions, both within tumors and across patients (11,12). The data suggest that this constitutes a fundamental dimension of cancer epigenomes, reflecting disrupted maintenance of epigenomic states and stochastic drift, which in turn may enable adaptation to the microenvironment, phenotypic plasticity, invasion, disease progression, and resistance to treatment (1,13,14). However, the genome-wide organization and functional consequences of DNA methylation variability across cancer types remain incompletely understood. Most prior studies have largely focused on individual cancer types, unpaired tumor samples, or isolated heterogeneity metrics, leaving open questions regarding the pan-cancer organization, regulatory context, and clinical relevance of DNA methylation variability (14–24).

Here, we address some of these open questions comprehensively through a systematic pan-cancer analysis of DNA methylation variability using paired tumor–normal samples across 16 cancer types. By integrating complementary measures of epigenetic variability, entropy, and stemness with regulatory annotation, transcription, and chromatin factor enrichment, genomic instability, and survival outcomes, we present a unified framework for understanding DNA methylation heterogeneity as a pervasive, biologically structured, and clinically consequential feature of tumor epigenomes.

## RESULTS

### DNA methylation variability is a pervasive feature of tumor epigenomes

To assess differences in DNA methylation variability between normal and tumor tissues, we analyzed paired Illumina 450K array data from The Cancer Genome Atlas (TCGA) across 16 cancer types (LUSC, n = 40; BLCA, n = 21; KIRP, n = 45; UCEC, n = 33; BRCA, n = 91; READ, n = 7; ESCA, n = 16; LUAD, n = 29; PAAD, n = 10; HNSC, n = 49; CHOL, n = 9; THCA, n = 56; LIHC, n = 50; KIRC, n = 158; PRAD, n = 50; COAD, n = 38; Total pairs, n = 702). CpG sites were ordered according to their median methylation levels in normal tissues in ascending order, from low to high methylation states. Tumor samples exhibited marked fluctuations in median methylation levels compared with matched normal tissues, indicative of increased variability. While the overall methylation landscape was generally preserved, tumor samples showed pronounced dispersion, particularly in intermediately methylated CpG sites, resulting in significantly wider interquartile ranges with higher data spread out across the majority of tumor types **(Figure 1A and B; Supplementary Figures 1 and 2)**, again illustrating higher variability. Consistent patterns of elevated methylation variability were also observed at the chromosomal level across all autosomes **(Figure 1C; Supplementary Figures 3)**. Taken together, these data support DNA methylation variability as a widespread feature observed across multiple tumor types.

**Figure 1.**
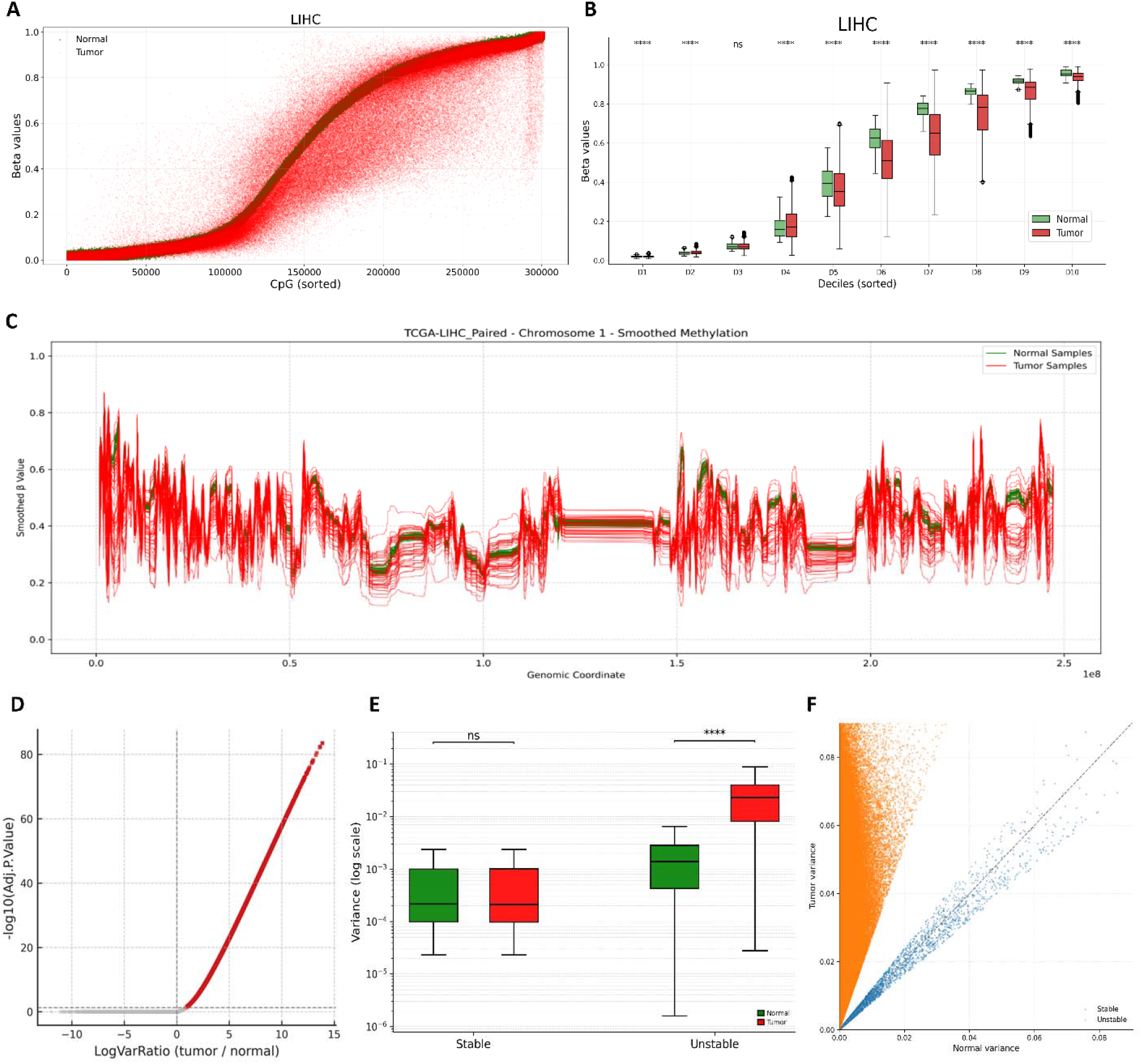
DNA methylation variability in paired tumor and normal samples from. LIHC is presented as one representative cancer type. Analyses were performed across all cancer types included in the study, as shown in Supplementary Figures 1-5. A) Scatter plot of DNA methylation β-values for individual CpG sites in tumor (red) and matched normal (green) samples, with CpGs sorted in ascending order by median methylation level in normal tissue. B) Boxplots showing β-value distributions in normal (green) and tumor (red) samples across deciles of CpG sites, with CpGs ranked by increasing median methylation in normal tissue. Tumor–normal differences within each decile were assessed using Mann–Whitney U tests with Benjamini–Hochberg false discovery rate (FDR) correction; significance is indicated by asterisks, C) Smoothed chromosome-level methylation profiles across Chromosome 1 for LIHC and normal samples, D) Volcano plot of log_2_ variance ratios (tumor/normal) versus −log_10_ adjusted P-values from differential variability testing, showing a large number of variable sites E) Boxplots showing methylation variance for CpG sites classified as stable or unstable, displayed separately for tumor and normal samples on a logarithmic scale showing that variability is observed exclusively in tumor samples and absent in matched normal tissues, and stable regions remained largely unchanged in both groups. Tumor–normal variance distributions within each class were compared using two-sided Mann–Whitney U tests on log_10_-transformed variance values, with statistical significance, F) Scatter plot of tumor versus normal variance per CpG site, with points colored by stability class and the identity line shown for reference.

To enable a more quantitative characterization of DNA methylation heterogeneity, we next sought to identify regions exhibiting differential variability between tumor and matched normal tissues within each cancer type **(Table 1)**. Differential variability analysis revealed a widespread increase in DNA methylation variance across most cancer types, driven by distinct subsets of CpG sites. In the majority of tumors, thousands of CpGs were classified as variably methylated (unstable), with an average unstable-to-stable ratio of 6.39 (range: 0.25–18.63) as shown in **Table 1**, demonstrating that increased DNA methylation variability of one group of CpG sites is a prominent and pervasive tumor-specific feature **(Figure 1D; Supplementary Figure 4)**. The terms “unstable” and “variable” are used interchangeably throughout the manuscript. Notably, this variability was observed exclusively in tumor samples and was absent in matched normal tissues, and stable regions remained largely unchanged in both groups **(Figure 1E and F; Supplementary Figure 5)**. These findings indicate the coexistence of stable and unstable DNA methylation CpG sites within tumor epigenomes.

**Table 1.** Summary of stable and unstable DNA methylation regions.

| **Cancer Type** | **Stable regions,**<br>**n** | **Unstable**<br>**regions, n** | **Unstable/stable**<br>**ratio** | **Median Var**<br>**ratio**<br>**(Stable)** | **Median**<br>**Log2 Var ratio**<br>**(Unstable)** |
| --- | --- | --- | --- | --- | --- |
| BLCA | 25 536 | 120 196 | 4.71 | 0.98 | 3.53 |
| BRCA | 23 719 | 191 831 | 8.09 | 0.99 | 3.04 |
| CHOL | 36 637 | 47 976 | 1.31 | 0.99 | 5.87 |
| COAD | 16 107 | 187 353 | 11.63 | 1.01 | 3.22 |
| ESCA | 50 466 | 29 045 | 0.58 | 1.01 | 4.56 |
| HNSC | 33 797 | 170 776 | 5.05 | 1.00 | 3.19 |
| KIRC | 21 189 | 239 373 | 11.30 | 1.03 | 2.94 |
| KIRP | 26 359 | 218 077 | 8.27 | 0.99 | 3.82 |
| LIHC | 16 004 | 206 758 | 12.92 | 1.00 | 3.68 |
| LUAD | 25 978 | 136 467 | 5.25 | 0.98 | 3.58 |
| LUSC | 11 771 | 219 298 | 18.63 | 1.01 | 3.60 |
| PAAD | 47 328 | 11 664 | 0.25 | 0.98 | 6.18 |
| PRAD | 36 649 | 73 588 | 2.01 | 1.00 | 2.53 |
| READ | 25 710 | 61 896 | 2.41 | 0.98 | 6.99 |
| THCA | 42 489 | 143 527 | 3.38 | 0.98 | 2.70 |
| UCEC | 35 652 | 121 324 | 3.40 | 0.98 | 3.54 |

### Variable DNA methylation regions are enriched in developmentally regulated chromatin states

To investigate whether stable and unstable methylation regions occupy distinct regulatory contexts, we performed locus overlap analysis using LOLA (25), leveraging publicly available regulatory and chromatin-associated annotations from the H1 human embryonic stem cell (H1 hESC) line, compiled from (26–28). We used the embryonic stem cell reference because it provides a unified baseline for assessing contexts associated with cellular identity, differentiation, and epigenomic reprogramming. Stably methylated regions were depleted in promoter regions and preferentially overlapped open sea CpG elements, SINE elements, and were enriched in transcribed chromatin and transcription-coupled and gene body-associated histone marks such as H3K4 methylations, and H3K36me3 **(Figure 2; Supplementary Figure 6)**.

**Figure 2.**
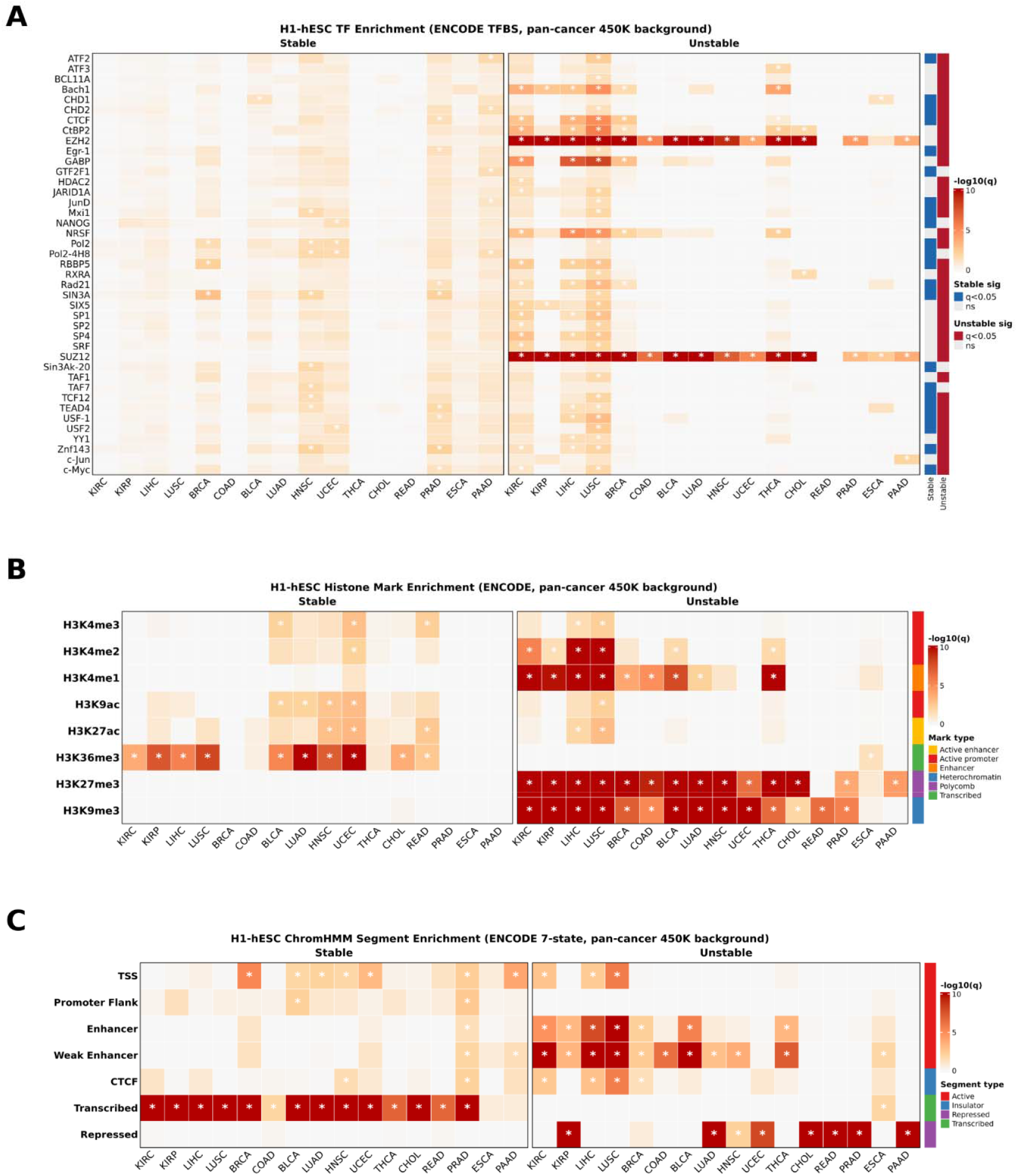
Regulatory and chromatin enrichment of stable and unstable DNA methylation regions across cancer types based on data from H1-hESC. A) Heatmap summarizing recurrent transcription factor (TF) enrichments across cancer types for stable and unstable regions, B) Heatmap summarizing recurrent histone modifications enrichments across cancer types for stable and unstable regions, C) Heatmap summarizing recurrent chromatin segment enrichments across cancer types for stable and unstable regions. This is related to Supplementary Figures 6-10.

In contrast, variably methylated or so-called unstable regions showed significant enrichment for binding sites of developmental and chromatin-repressive transcription factors, ostensibly Polycomb-associated regulators EZH2 and SUZ12, the biogenesis, development, and cell cycle and cell proliferation factor, GABP (also known as NRF2), the developmental repressor NRSF, as well as the major architectural protein CTCF, and the chromatin remodeler CHD1. These regions were further enriched for satellite DNA, repressive histone modifications such as H3K27me3 and H3K9me3, alongside dynamically regulated enhancer histone marks such as H3K4me1/2, a combination characteristic of facultative and developmentally poised regulatory regions **(Figure 2; Supplementary Figure 6)**. In addition, analysis of ATAC-seq data from TCGA revealed that stable regions exhibit higher chromatin accessibility than unstable regions, whereas unstably methylated regions showed reduced accessibility consistent with heterochromatin and weakly transcribed chromatin states **(Supplementary Figure 7)**. Together, these findings indicate that DNA methylation variability in cancer likely reflects instability of developmentally regulated chromatin states rather than stochastic epigenetic noise.

To validate these observations under more conservative criteria, we next defined pan-stable and pan-unstable DNA methylation regions across cancer types. Regions were designated pan-stable if they were called stable in ≥6 of 16 cancer types and were never called unstable in any cancer type, whereas pan-unstable regions were defined as those called unstable in ≥13 of 16 cancer types and never called stable in any cancer type; all classifications were restricted to autosomal loci **(Supplementary Figure 8)**. Owing to the limited abundance of pan-stable regions, robust statistical analysis was not feasible, albeit preliminary observations showed enrichment in H3K36me3 and transcribed regions. In addition, consistent with our earlier findings, pan-unstable regions were enriched in EZH2 and SUZ12 binding sites, while chromatin state analysis further showed enrichment in H3K9me3 and H3K27me3-decorated heterochromatin and repressed segments, with depletion from active chromatin states **(Supplementary Figure 9)**. Finally, transcription factor motif enrichment analysis of pan-unstable regions using HOMER (29) followed by Gene Ontology analysis (30) revealed significant associations with biological processes related to cell fate determination, differentiation and development, tissue remodeling and signaling, as well as stress and environmental response pathways **(Supplementary Figure 9D)**. In addition, CpG sites classified as pan-unstable exhibited consistently higher DNA methylation variability across multiple cancer types compared with pan-stable sites **(Supplementary Figure 10)**. Collectively, these results demonstrate that increased DNA methylation variability in cancer arises within developmentally regulated and epigenetically plastic genomic regions.

### DNA methylation disorder is more prevalent in tumors than in normal tissues

Because DNA methylation variability arises in non-random, epigenetically relevant genomic regions and varies between tumors from different patients, we next sought to evaluate its biomedical significance in a framework using standardized, intra-tumor quantitative measures of methylation disorder and stemness properties. To this end, we calculated three complementary metrics: the proportion of intermediately methylated sites (PIM), genome-wide Shannon entropy (SE), and a DNA methylation– based stemness index (mDNAsi or SI), hereafter collectively referred to as measures of DNA methylation disorder/entropy, although the stemness index might be a related but not identical epigenetic dimension (16,31,32).

The distribution of DNA methylation disorder in tumors was right-shifted relative to matched normal tissues, meaning that they consistently exhibit higher DNA methylation disorder than normal tissues, regardless of how it is quantified **(Figure 3A, B, and C; Supplementary Figure 11A, B, and C)**. Elevated methylation disorder was observed across the majority of cancer types, although the magnitude of this effect varied by cancer type, highlighting cancer-specific patterns of methylome instability **(Supplementary Figure 21D-I)**. Importantly, differences in methylation disorder were not driven by cohort size but were reproducible within cancer types and consistent across the pan-cancer dataset **(Supplementary Figure 21G, H, and I)**.

**Figure 3.**
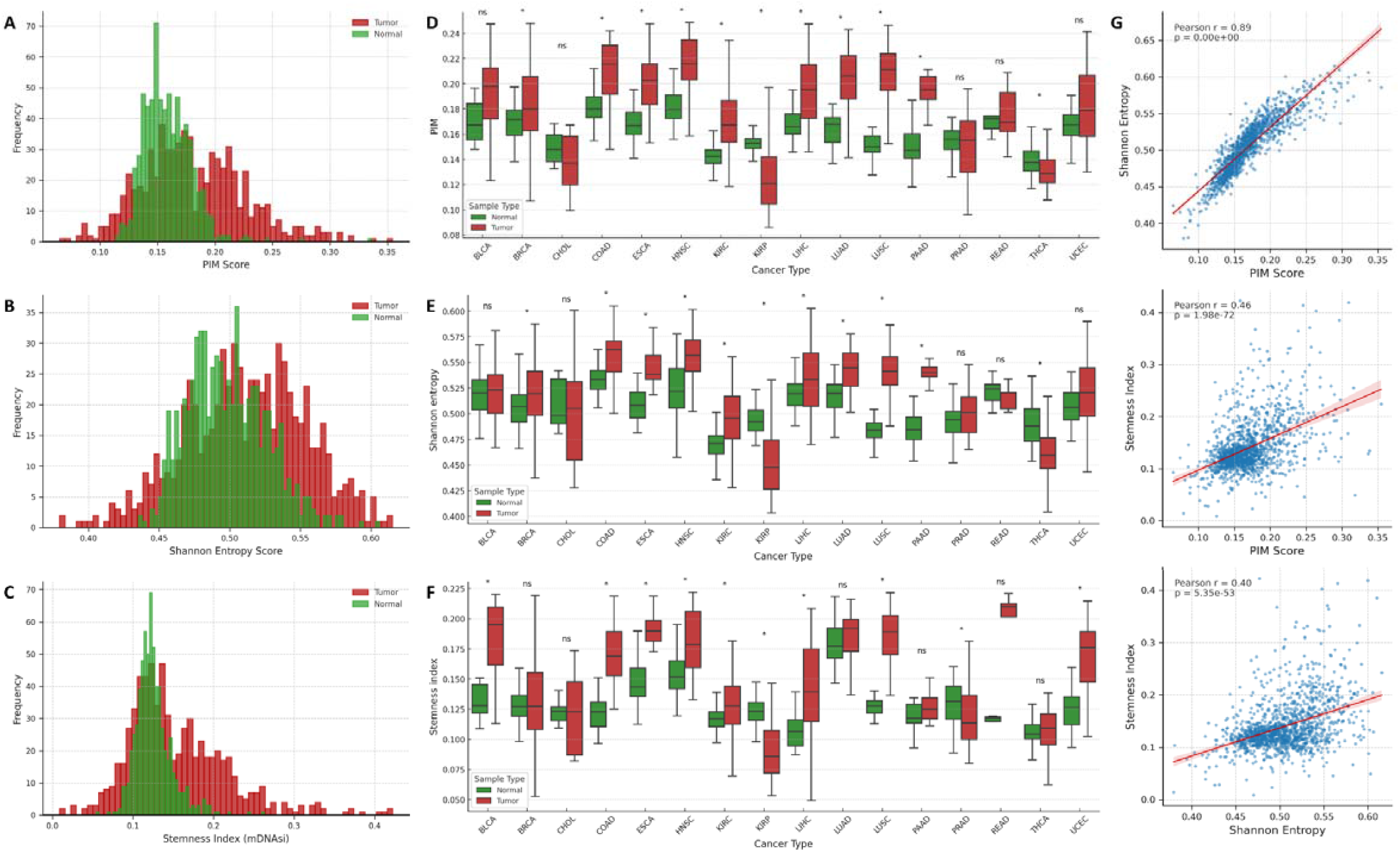
Genome-wide DNA methylation disorder across cancer types quantified by Proportion of Intermediately Methylated sites (PIM), Shannon entrop, and methylation-derived stemness (mDNAsi). Overlaid histograms of the A) PIM, B) Shannon entropy, and C) Stemness index - mDNAsi for tumor (red) and matched normal (green) samples across all cancers. Boxplots comparing D) PIM, E) Shannon entropy, and F) Stemness index - mDNAsi for tumor (red) and matched normal (green) samples per cancer type. Statistical significance was calculated with the Wilcoxon paired test. G) Correlation scatter plots of PIM, Shannon entropy, and mDNAsi with linear regression lines shown for reference and Pearson correlation coefficients reported. This is related to Supplementary Figure 11.

Correlation analyses revealed a strong association between PIM and Shannon entropy (Pearson’s r = 0.89), indicating that a substantial component of global methylation disorder emanates from intermediately methylated CpG sites. In contrast, both PIM and Shannon entropy showed only moderate correlations with mDNAsi (Pearson’s r = 0.46 and r = 0.40, respectively), suggesting that these metrics capture overlapping but non-identical methylation dimensions **(Figure 3G)**. The consistency of these patterns across cancer types indicates that methylation disorder represents a pervasive feature of tumor methylomes.

### Elevated DNA methylation variability is associated with increased genomic instability and invasive tumor properties

Next, we investigated the association between DNA methylation disorder and a range of orthogonal measures of genomic instability and clinical characteristics. For each methylation disorder metric, such as PIM, Shannon entropy (labeled as SE), and mDNAsi (labeled as SI), we stratified the pool of tumors into five groups in ascending order from low to high disorder. Notably, samples assigned to corresponding quintiles across the three metrics showed substantial overlap; tumor types such as non– small cell lung cancer, head and neck cancer, hepatobiliary cancers, and colorectal cancer were overrepresented in high-disorder groups, whereas renal and thyroid cancers were depleted **(Supplementary Figure 12A and B)**.

Across quintiles of PIM, Shannon entropy, and mDNAsi, higher methylation disorder was accompanied by stepwise increases in genomic features such as aneuploidy, tumor break load, fraction of genome altered, and tumor mutational burden **(Figure 4)**. Tumors assigned to corresponding strata across methylation disorder metrics exhibited concordant mutational landscapes, with specific genetic alterations preferentially associated with either high or low levels of methylation disorder (**Supplementary Figure 12C)**. These relationships were consistent across all three metrics. In a similar vein, tumors with higher disorder scores showed an increased likelihood of lymph node metastasis at presentation and were more prone to recurrence following initial therapy **(Figure 5A and B)**. Along the same lines, Ragnum scores increased stepwise with higher methylation disorder across all three metrics **(Figure 5C)**. The Ragnum hypoxia score is a transcriptome-based metric that quantifies tumor hypoxia by summarizing the expression of hypoxia-responsive gene programs, with higher scores indicating increased hypoxic signaling associated with more aggressive tumor biology (33). These trends remained consistent after normalization for tumor purity **(Supplementary Figures 13 and 14)**. In summary, DNA methylation variability not only reflects epigenomic but also genomic instability and marks certain tumor types, with increased invasive potential, recurrence risk, and aggressive clinical behavior.

**Figure 4.**
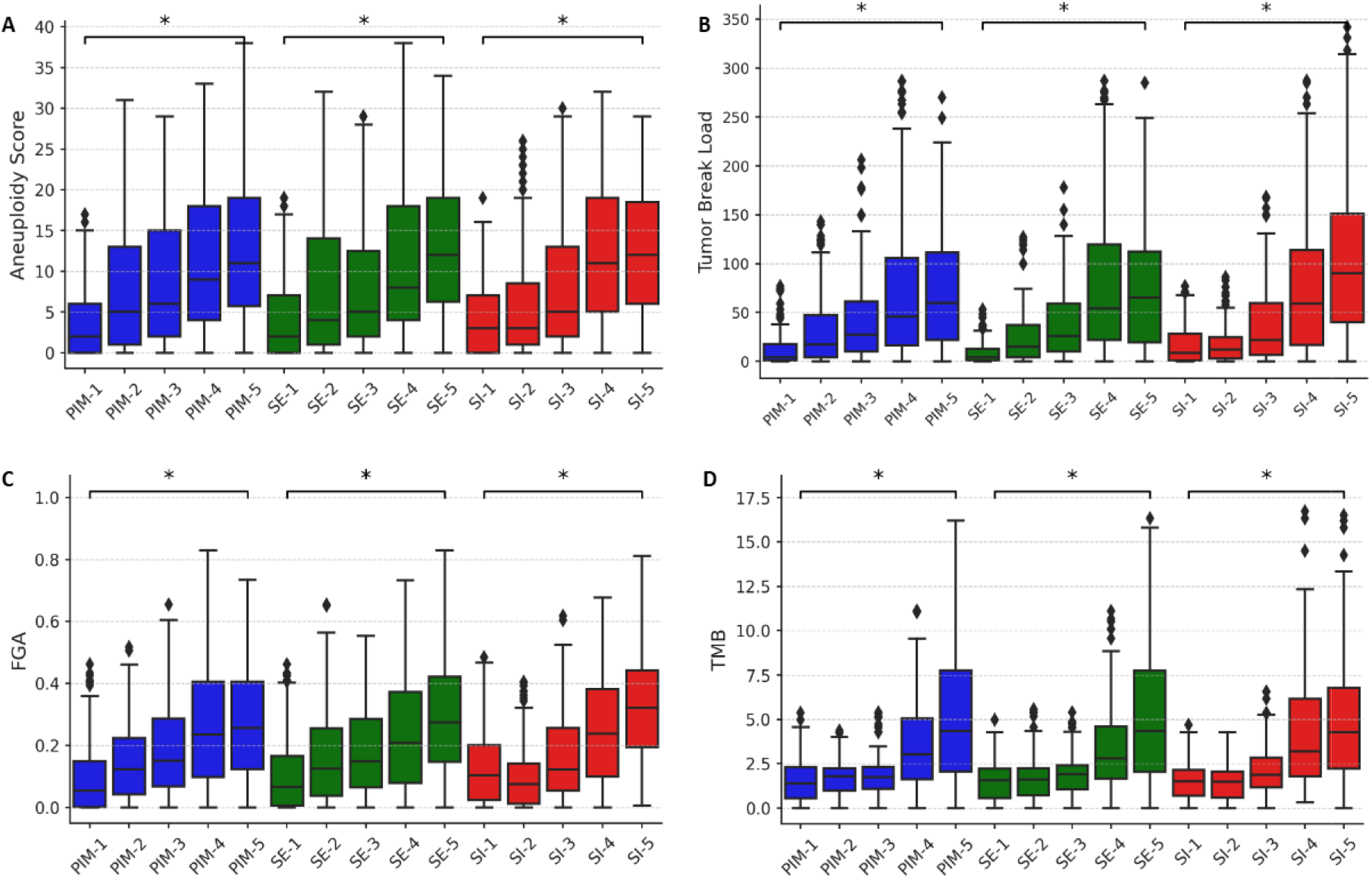
Association between DNA methylation disorder metrics and genomic instability features. Boxplots of tumors stratified into quintiles (1–5) based on increasing levels of DNA methylation disorder as quantified by the Proportion of Intermediately Methylated sites (PIM; blue), Shannon entropy (SE; green), and methylation-derived stemness index (SI/mDNAsi; red). Boxplots show the distribution of genomic instability measures across quintiles for A) aneuploidy score, B) tumor break load, C) fraction of genome altered (FGA), and D) tumor mutational burde (TMB). Boxes represent the interquartile range with median values indicated; whiskers denote 1.5× IQR, and points represent outliers. Brackets indicate comparisons across increasing quintiles within each metric, with statistical significance calculated using the Mann–Whitney U test. The signals are unadjusted for tumor purity. This is related to Supplementary Figures 12 and 13.

**Figure 5.**
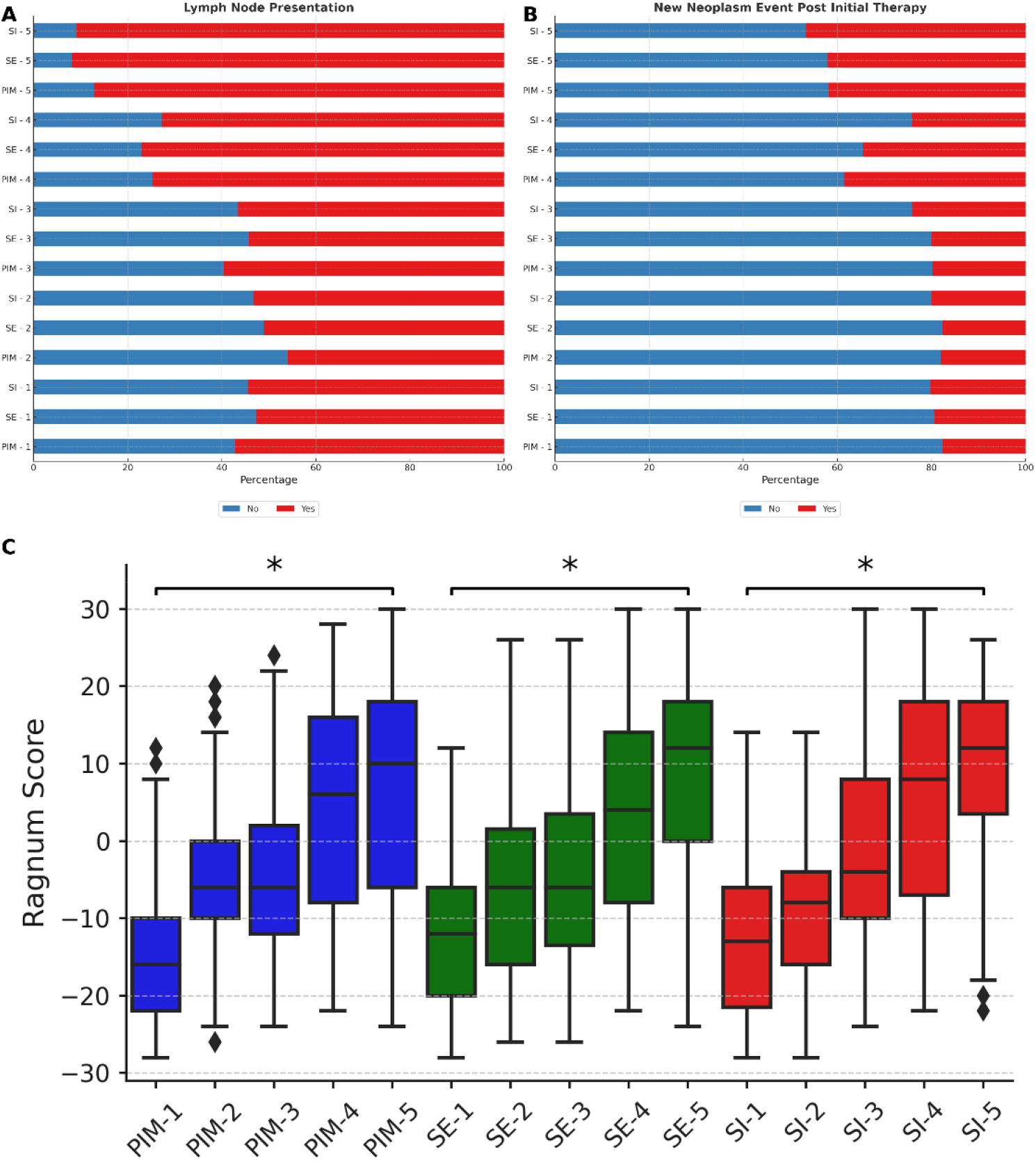
Clinical associations of DNA methylation disorder with metastatic presentation, recurrence, and tumor hypoxia. Stacked bar plots summarizing the proportion of tumors with A) lymph node involvement at diagnosis and B) new neoplasm events following initial therapy, stratified by quintiles (1–5) of DNA methylation disorder defined by PIM, Shannon entropy (SE), and methylation-derived stemness index - mDNAsi (SI). Bars indicate the percentage of cases classified as “Yes” or “No” for each clinical feature within each quintile. C) Boxplots of the Ragnum hypoxia score across the same disorder quintiles for PIM (PIM, blue), Shannon entropy (SE, green), and mDNAsi (SI, red), unadjusted for tumor purity. Boxes represent the interquartile range with median values indicated; whiskers denote 1.5× IQR, and points represent outliers. Brackets indicate comparisons across increasing quintiles within each metric, with statistical significance calculated using the Mann–Whitney U test. This is related to Supplementary Figures 12 and 14.

### Elevated DNA methylation variability is associated with adverse clinical outcomes and reduced patient survival

Furthermore, we examined the association between DNA methylation disorder and patient survival. Stratification of tumors by increasing levels of methylation disorder revealed a strong, level-dependent relationship with adverse clinical outcomes. Across quintiles of PIM, Shannon entropy, and mDNAsi, higher methylation disorder was consistently associated with progressively worse overall, progression-free, and disease-free survival, with clear separation of survival curves and hazard risks **(Figure 6)**. Relative to the lowest-disorder group, patients in the highest disorder quintile exhibited a substantially increased risk of death and disease recurrence, with hazard ratios ranging from approximately 2.2-to 2.7-fold for overall survival, 4- to 13-fold for disease-free survival, and 2- to 2.5-fold for progression-free survival, depending on the disorder metric. These trends in survival per methylation disorder metric and per stage remained consistent after normalization for tumor purity (**Supplementary Figures 15 and 16)**. Together, these findings indicate that increasing DNA methylation disorder is strongly associated with adverse clinical outcomes, supporting methylome instability as a robust marker of tumor aggressiveness and poor prognosis.

**Figure 6.**
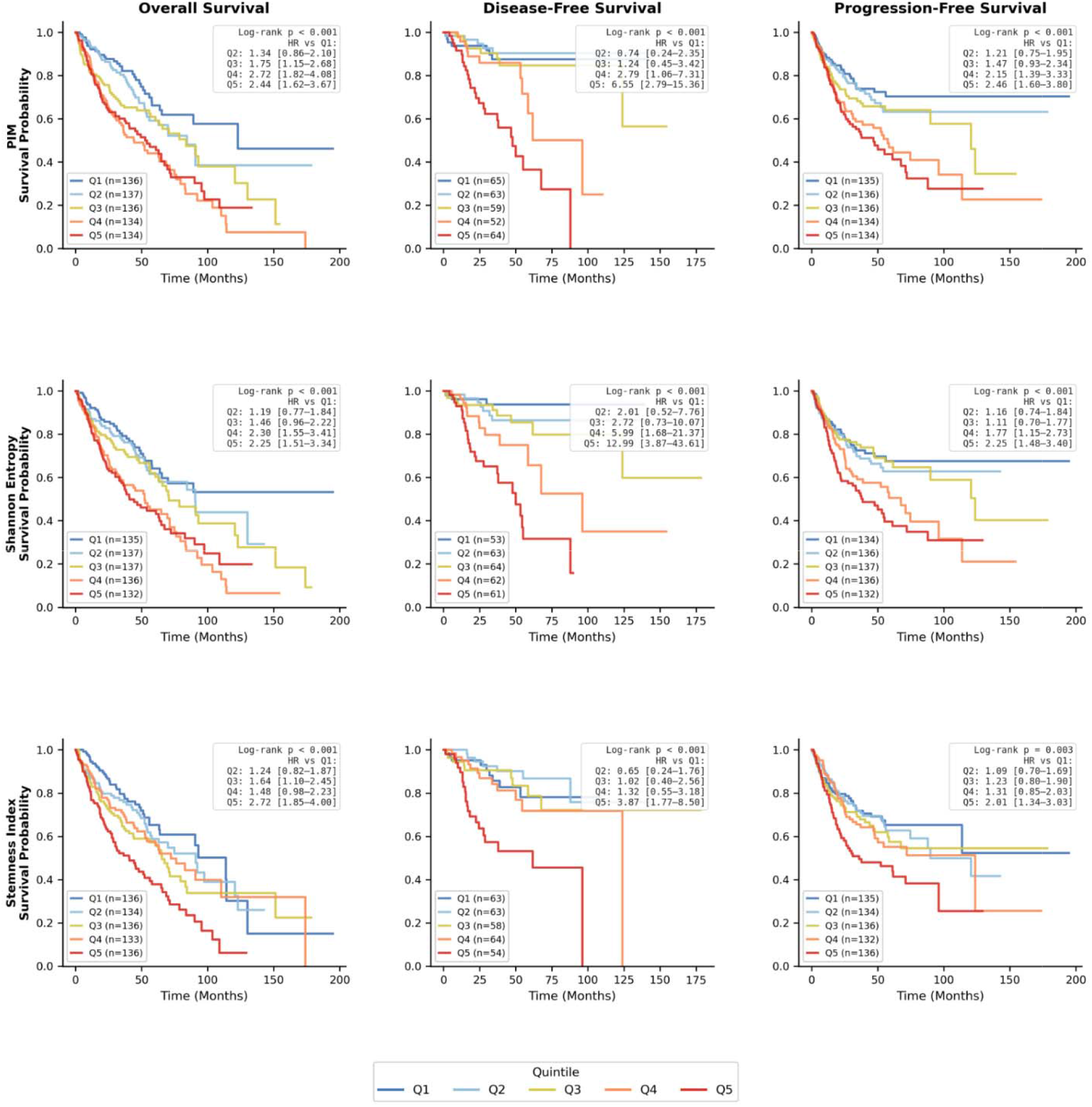
Survival analyses stratified by DNA methylation disorder metrics. Kaplan–Meier survival curves are shown for overall survival (left), disease-free survival (middle), and progression-free survival (right) unadjusted for tumor purity, with patients stratified into quintiles (1–5) according to increasing levels of DNA methylation disorder as quantified by the proportion of intermediately methylated sites (PIM), Shannon entropy, and the methylation-derived stemness index mDNAsi. Curves are color-coded by quintiles. Log-rank test P values for each comparison are indicated within panels. Hazard ratios with 95% confidence intervals for each quintile relative to the lowest-disorder group are reported in the legends of the respective panels. This is related to Supplementary Figures 15 and 16.

## DISCUSSION

In this study, we conducted a systematic pan-cancer analysis of DNA methylation variability using paired tumor–normal samples across 16 cancer types. Our results indicate that increased DNA methylation variability is a widespread feature of tumor epigenomes. Tumor samples consistently exhibited increased dispersion of methylation values, particularly at intermediately methylated CpG sites. These findings support the notion that altered regulation of DNA methylation variability represents a general property of tumors rather than a tumor-type–specific phenomenon.

We further show that DNA methylation variability is non-randomly distributed and localizes to specific regulatory and chromatin contexts. Variably methylated regions were enriched in enhancer compartments, chromatin states associated with heterochromatin, weak transcription, and Polycomb-mediated repression, whilst they were depleted in regions of active transcription. Enrichment for developmental and chromatin-repressive transcription factor binding sites suggests that DNA methylation variability may be associated with perturbations of developmental regulatory programs.

At the global level, we demonstrate that DNA methylation disorder quantified using complementary metrics, including PIM, Shannon entropy, and a DNA methylation–based stemness index, is consistently elevated in tumors compared with normal tissues. Higher levels of methylation disorder were associated with multiple orthogonal measures of genomic instability, including aneuploidy, fraction of genome altered, tumor break load, and tumor mutational burden. In addition, methylation disorder correlated with clinically aggressive features such as lymph node involvement, post-therapy recurrence, hypoxia-associated transcriptional programs, and reduced stage-dependent overall, progression-free, and disease-free survival. These associations were observed across independent disorder metrics and exhibited level-dependent trends, indicating that methylation disorder captures biologically and clinically relevant aspects of tumor behavior, without implying causality.

Increased DNA methylation variability may reflect impaired propagation of epigenetic states, particularly within developmentally regulated and facultative chromatin domains, leading to progressive diversification of regulatory programs or regulatory potentials within tumors. Such epigenetic drift could increase phenotypic heterogeneity, relax lineage constraints, and expand the range of transcriptional states accessible to cancer cells, thereby facilitating adaptation to hypoxic, immune, or therapeutic pressures. In this context, it stands to reason that elevated methylation variability, as shown in our findings, would associate with more aggressive behavior and poorer clinical outcomes, not as a direct driver but as a marker of increased epigenomic instability and evolutionary potential within tumors. In line with this interpretation, cancer types that are generally associated with less favorable clinical outcomes tended to show higher levels of DNA methylation variability, whereas tumor types with more favorable prognoses tended to exhibit lower levels, although substantial heterogeneity exists within and across cancer types.

Several limitations should be taken into account. First, our analyses are based on bulk DNA methylation profiles and therefore do not resolve cell-to-cell heterogeneity or fully account for tumor purity, although we normalized for tumor purity, precise estimation of purity from bulk data remains technically and conceptually challenging. The use of paired tumor–normal samples and multiple orthogonal metrics mitigates, albeit not fully, some of these concerns. As a consequence, the observed variability likely reflects a combination of intra-tumor heterogeneity and inter-tumor differences across patients, which cannot be fully disentangled in bulk data. Second, the Illumina 450K array provides limited coverage of repetitive elements and large heterochromatic regions, potentially underestimating variability in these compartments. Third, while embryonic stem cell–derived chromatin annotations provided a unified reference for assessing developmental and identity-related regulatory contexts, tissue-specific chromatin landscapes may reveal additional layers of variability. Finally, the observational nature of this study precludes causal inference, and functional and single-cell approaches will be required to determine whether DNA methylation variability actively contributes to tumor progression or primarily reflects downstream consequences of oncogenic stress.

## CONCLUSION

Overall, our findings identify increased DNA methylation variability as a pervasive feature of tumor epigenomes that is reproducibly associated with genomic instability and adverse clinical outcomes across cancers. These results suggest that methylome disorder captures an important dimension of tumor biology reflecting epigenomic instability and expanded regulatory plasticity. Future studies integrating single-cell epigenomic profiling and functional perturbation approaches will be necessary to determine how methylation variability arises and whether it can be leveraged for prognostic or therapeutic applications.

## MATERIALS AND METHODS

### Datasets used in the study

Publicly available molecular and clinical data were obtained from The Cancer Genome Atlas (TCGA) consortium, a large-scale cancer multi-omics initiative that has molecularly characterized over 20,000 primary tumors and matched normal samples across 33 cancer types. We analyzed paired tumor– normal samples from 16 cancer types for which matched data were available. Datasets were accessed through the cBioPortal for Cancer Genomics (https://www.cbioportal.org) and the Genomic Data Commons (GDC) data portal (https://portal.gdc.cancer.gov/).

Genome-wide DNA methylation profiles were generated using the Illumina HumanMethylation450 (450K) BeadChip platform, which interrogates over 450,000 CpG sites spanning promoter regions, gene bodies, CpG islands, shores, shelves, and open-sea regions. Level 3 beta values, representing the proportion of methylated alleles at each CpG site and ranging from 0 (unmethylated) to 1 (fully methylated), were used for all analyses. CpG probes were filtered to exclude those with missing values or mapping to sex chromosomes, where appropriate. Paired tumor–normal methylation profiles were analyzed to assess differences in DNA methylation variability and disorder while controlling for inter-individual baseline methylation patterns. DNA methylation datasets were retrieved via the GDC application programming interface (API), assembled into matched tumor–normal matrices on a high-performance computing cluster, and used for subsequent analyses. The Biomni platform was used to validate some of the observations (34).

Somatic mutation and copy number alteration, clinical annotations including measures of genomic instability, lymph node involvement, cancer recurrence, survival outcomes, and pathological features were retrieved from TCGA clinical metadata deposited in cBioPortal. Chromatin accessibility data were obtained from the GDC portal in the form of TCGA ATAC-seq bigWig files, where available, and were used to evaluate associations between DNA methylation stability and chromatin accessibility by overlapping CpG coordinates with accessible chromatin regions.

### DNA methylation variability analyses

#### Stable and unstable (variable) regions

Differential variability analysis was performed using the DiffVar function implemented in the missMethyl Bioconductor package (35,36), which applies a limma-based empirical Bayes framework for testing differences in variance between paired tumor and normal samples. For each CpG site, differential variability between tumor and normal samples was tested using an empirical Bayes–moderated F-test for equality of variances, accounting for the paired study design. Tumor-associated changes in variability were summarized using the log_2_ variance ratio (log_2_[Var_tumor /Var_normal]). Resulting p-values were adjusted for multiple testing using the Benjamini–Hochberg false discovery rate (FDR). CpG sites were classified as stable when variability was comparable between groups (0.8 ≤ Var_tumor /Var_normal ≤ 1.2 and FDR > 0.05) and as unstable when variability was significantly increased in tumors (log_2_ variance ratio > 1 and FDR ≤ 0.001); all other CpGs were excluded from stability-based analyses.

For visualization of genome-wide patterns, chromosomal methylation profiles were smoothed by aggregating CpG-level beta values across genomic windows to reduce local noise and highlight large-scale variability trends.

Variance estimates were summarized by cancer type, sample group, and stability class and visualized using log-scaled boxplots with outliers suppressed. Tumor–normal differences were assessed descriptively using two-sided Mann–Whitney U tests on log-transformed variance values. Tumor versus normal variance scatter plots were generated with an identity line and high-percentile axis limits to highlight tumor-specific variance inflation.

#### Pan stable and pan unstable regions

For each TCGA cancer type, DiffVar results were converted into hg19 genomic intervals by expanding each CpG to a fixed ±125 bp window and merging adjacent windows separated by ≤50 bp to generate non-redundant regions. Per-cancer stable and unstable region sets were then combined across all cancer types to create a global interval catalogue. Each interval in this catalogue was intersected with the per-cancer region sets to record, for every interval, the number of cancer types in which it was classified as stable or unstable. Cross-cancer consistency was evaluated by varying a support parameter k (k = 1–16), defined as the minimum number of cancer types required to support stability or instability for a given interval. Pan-cancer stable and unstable regions were defined using a strict criterion,≥k, supporting cancer types with no opposing classifications.

Pairwise similarity of stable and unstable methylation-variability regions across cancer types was quantified using the Jaccard index computed on autosomal BED intervals. For each TCGA cancer type, probe-centered regions were standardized to fixed 250 bp windows, merged to remove fragmentation, and restricted to autosomes. Similarity between each pair of cancer types was computed separately for stable and unstable region sets using bedtools jaccard, which measures length-weighted base-pair overlap. Pairwise Jaccard values were assembled into symmetric matrices and visualized as heatmaps. This helped us in the definition of thresholds used to classify pan-stable and pan-unstable regions.

#### Genomic and chromatin state annotations

Regulatory enrichment analyses were performed using LOLA (Locus Overlap Analysis) (25). BED-formatted CpG-centered genomic windows representing stable or unstable DNA methylation regions per cancer type were used as query sets, with the full set of eligible Illumina 450K CpG windows used as the background universe. Enrichment was evaluated against publicly available ChIP-seq peak annotations from the H1 human embryonic stem cell (H1-hESC) context, including ENCODE transcription factor binding sites and Roadmap Epigenomics histone modification datasets, accessed via the LOLA RegionDB (hg19). The same was done for chromatin-state annotation using the hg19 ChromHMM segmentations (37) taken from H1-hESC. Chromatin-state distributions were summarized relative to a 450K-matched background to assess enrichment or depletion of unstable regions across major chromatin states.

Overlap enrichment was quantified using odds ratios, q-values, and p-values, with multiple-testing correction applied where appropriate, and enrichment signals were summarized across cancer types for downstream analyses. The enrichment metric was selected based on the clarity and interpretability of the resulting visualizations.

Genomic annotation of stable and unstable DNA methylation variability regions was performed using pyBEDtools (38) BED-formatted CpG-centered regions representing stable, unstable, and background (Illumina HumanMethylation450K-derived) sets were annotated against the hg19 genome assembly UCSC genome annotation database for genic and CpG elements.

### Gene ontology

Gene Ontology (GO) over-representation analysis was performed to characterize biological programs associated with pan-cancer unstable DNA methylation variability regions. Motif enrichment was first conducted on unstable-region BED intervals using HOMER (29) with an Illumina HumanMethylation450K-derived, design-matched genomic background to control for array coverage bias. Enriched motifs were consolidated to transcription factor (TF)–level candidates by selecting the most significant motif per TF and applying significance thresholds (P < 0.05, fold enrichment ≥ 1.5). TFs were then mapped to curated TF–target gene sets (Enrichr ChEA_2016 and ENCODE_TF_ChIP-seq_2015) (39), and the resulting gene set was intersected with a 450K-matched gene universe. GO enrichment analysis was performed using clusterProfiler (40) with Benjamini–Hochberg correction, enriched Biological Process terms were visualized using dot plots.

### ATAC-seq

Normalized density signals were summarized across DNA methylation–defined stable and unstable region sets by aggregating ATAC-seq signal within ±2 kb windows centered on each region, enabling comparison of chromatin accessibility patterns (two-sided Mann-Whitney U test) between region classes across cancer types.

### Methylation disorder metrics: PIM, Shannon entropy, and stemness index

Genome-wide intermediate methylation was quantified using the Proportion of Intermediate Methylation (PIM). Illumina HumanMethylation450K β-values were used as provided from the preprocessed β-value matrix. For each sample, CpG sites were classified as intermediately methylated if their β-values fell within the range 0.2 ≤ β < 0.6, and PIM was calculated as the number of CpGs in this intermediate range divided by the total number of retained CpGs for that sample. Tumor–normal differences in PIM were visualized using overlaid histograms and cancer-stratified boxplots, and statistical significance was assessed using two-sided Mann–Whitney U tests for unpaired comparisons and paired Wilcoxon signed-rank tests for matched samples.

Global DNA methylation disorder was quantified at the sample level using Shannon entropy (41). After standard preprocessing of Illumina HumanMethylation450K arrays, β-values were treated as Bernoulli parameters for each CpG site and sample. To avoid numerical artefacts, β-values were constrained to the interval, [ε, 1 − ε] (ε = 1 × 10^−6^). CpG-level entropy was computed as

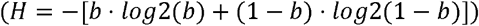

and a single per-sample entropy score was calculated as the mean entropy across all retained CpGs.

The methylation-derived stemness index (mDNAsi or SI) was obtained for each matched normal and primary tumor sample across TCGA cancer types from the original publication (31), and samples lacking mDNAsi values were excluded from subsequent analyses. To account for potential confounding by tumor purity (42) methylation disorder metrics were normalized by extracting the residuals from a linear regression model of the metric against tumor Consensus Purity Estimate (CPE), subsequently rescaled by the global mean.

Survival analysis was conducted in lifelines (43) and Kaplan-Meier curves were generated to estimate the probability of clinical events over time, and differences between patient groups were assessed using the log-rank test. Hazard ratios (HR) and their 95% confidence intervals (CI) were derived from the log-rank test statistics to quantify the relative risk between groups. A p-value of < 0.05 was considered statistically significant. Exploratory analyses of tumor type distribution and mutation frequencies across DNA methylation disorder groups were performed using the cBioPortal platform and its integrated visualization and summary tools.

### Statistics and visualizations

Genomic interval processing and visualization for this analysis were implemented in Python using bedtools, pandas, NumPy, Matplotlib, and SciPy. Selected calculations and visualizations were reproduced on the Biomni platform.

## Declarations

### Ethics approval and consent to participate

Not applicable. All data are taken from publicly available datasets and cancer projects.

### Consent for publication

Not applicable. All data are taken from publicly available datasets and already reported in the literature.

### Availability of data and materials

All data are publicly available at the cBioportal repository, https://www.cbioportal.org/, and the Genomic Data Commons data portal https://portal.gdc.cancer.gov/.

### Funding

Funded by the European Union under Horizon Europe (project ChatMED Grant Agreement ID: 101159214).

### Use of AI

ChatGPT (OpenAI) and Grammarly were utilized for language editing and stylistic improvement. No AI tools were used for data analysis, data interpretation, or the generation of scientific conclusions. All content was reviewed and approved by the authors.

### Conflict of interest

The authors declare no conflicts of interest, financial or otherwise, related to this work. Although some authors are employed by private companies, these affiliations did not influence the design, data collection, analysis, interpretation, or manuscript preparation. All authors maintained full objectivity, and the research was conducted independently.

### Author contributions

GK conceived, designed, and led the study. MSM supervised aspects of the computational and statistical analyses. DjB collected, curated, processed, and analyzed the data with the help of GK and BGj. GK wrote the initial draft of the manuscript. All authors contributed to data interpretation, critically revised the manuscript for important intellectual content, and approved the final version.

## Notes

https://www.cbioportal.org/

https://portal.gdc.cancer.gov/

### Competing Interest Statement

The authors have declared no competing interest.

