## Supplementary Figures for "DNA methylation variability defines a fundamental dimension of tumor epigenomes linked to genomic instability, tumor aggressiveness, and clinical outcomes"

**Corresponding author*

### Supplementary Figures

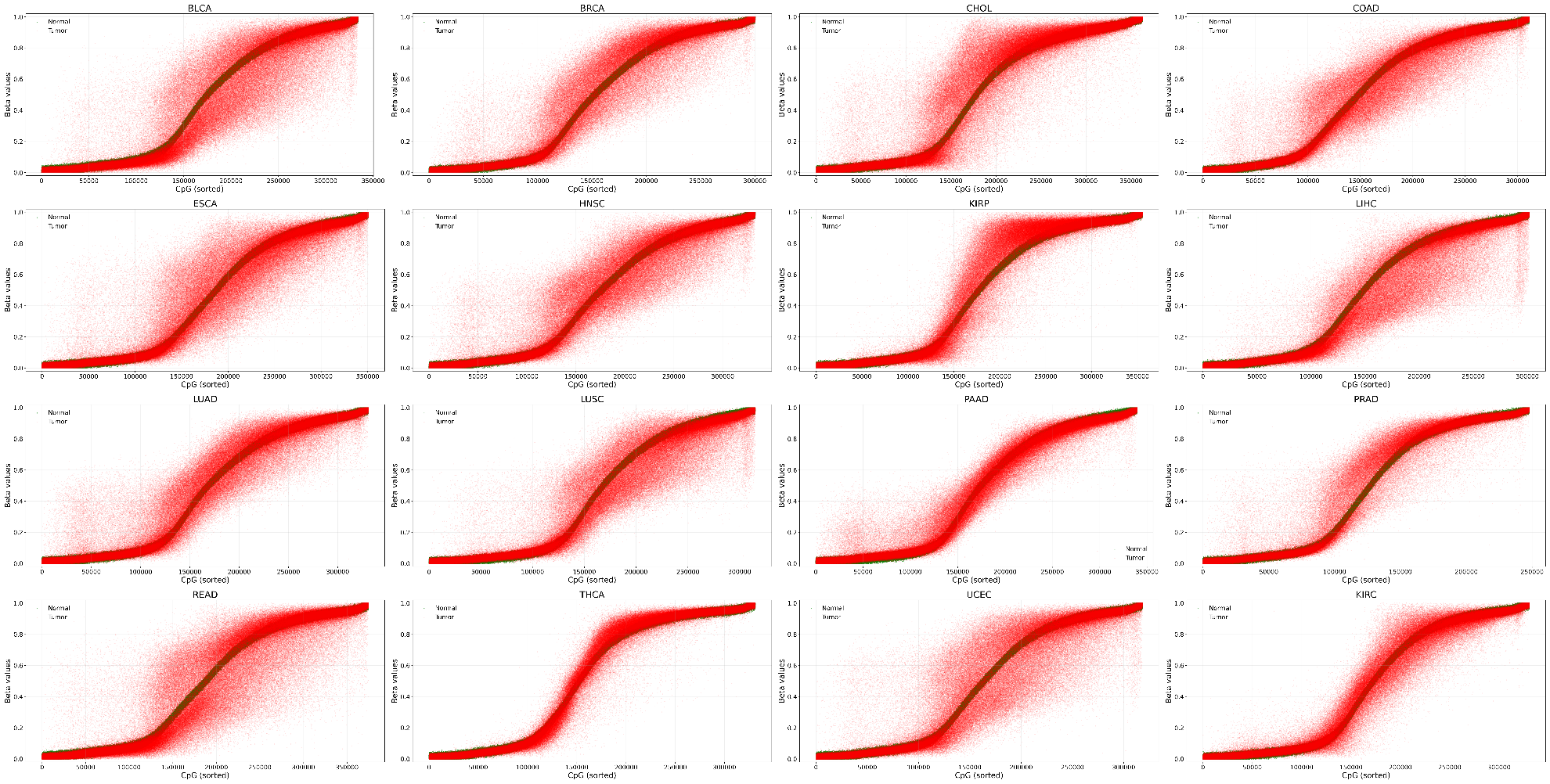

**Supplementary Figure 1. DNA methylation variability in paired tumor and normal samples from 16 types of cancer.** Scatter plot of DNA methylation β-values for individual CpG sites in tumor (red) and matched normal (green) samples, with CpGs sorted in ascending order by median methylation level in normal tissue.

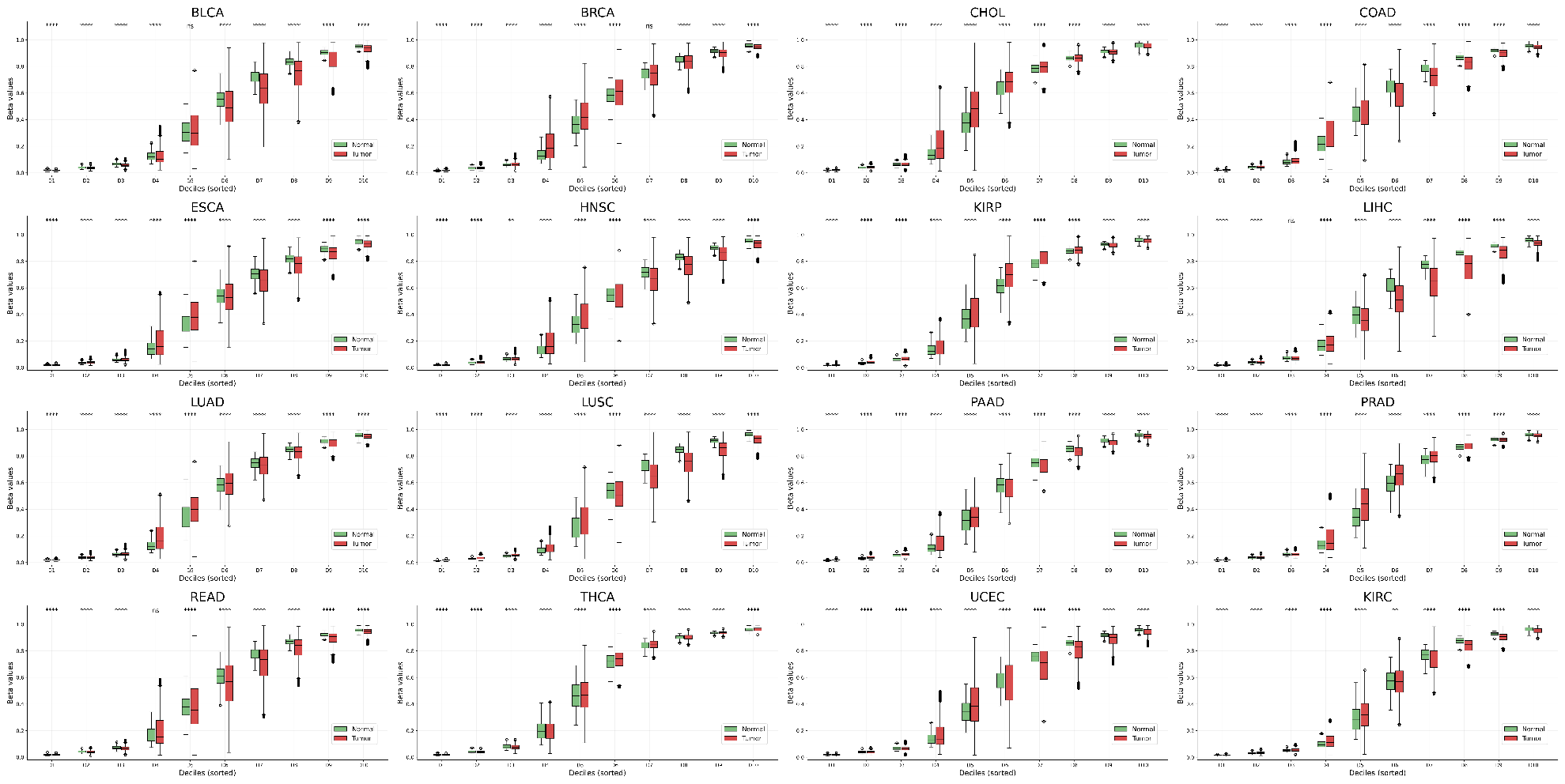

**Supplementary Figure 2. DNA methylation variability in paired tumor and normal samples from 16 types of cancer.** Boxplots showing β-value distributions in normal (green) and tumor (red) samples across deciles of CpG sites, with CpGs ranked by increasing median methylation in normal tissue. Tumor–normal differences within each decile were assessed using Mann–Whitney U tests with Benjamini–Hochberg false discovery rate (FDR) correction; significance is indicated by asterisks.

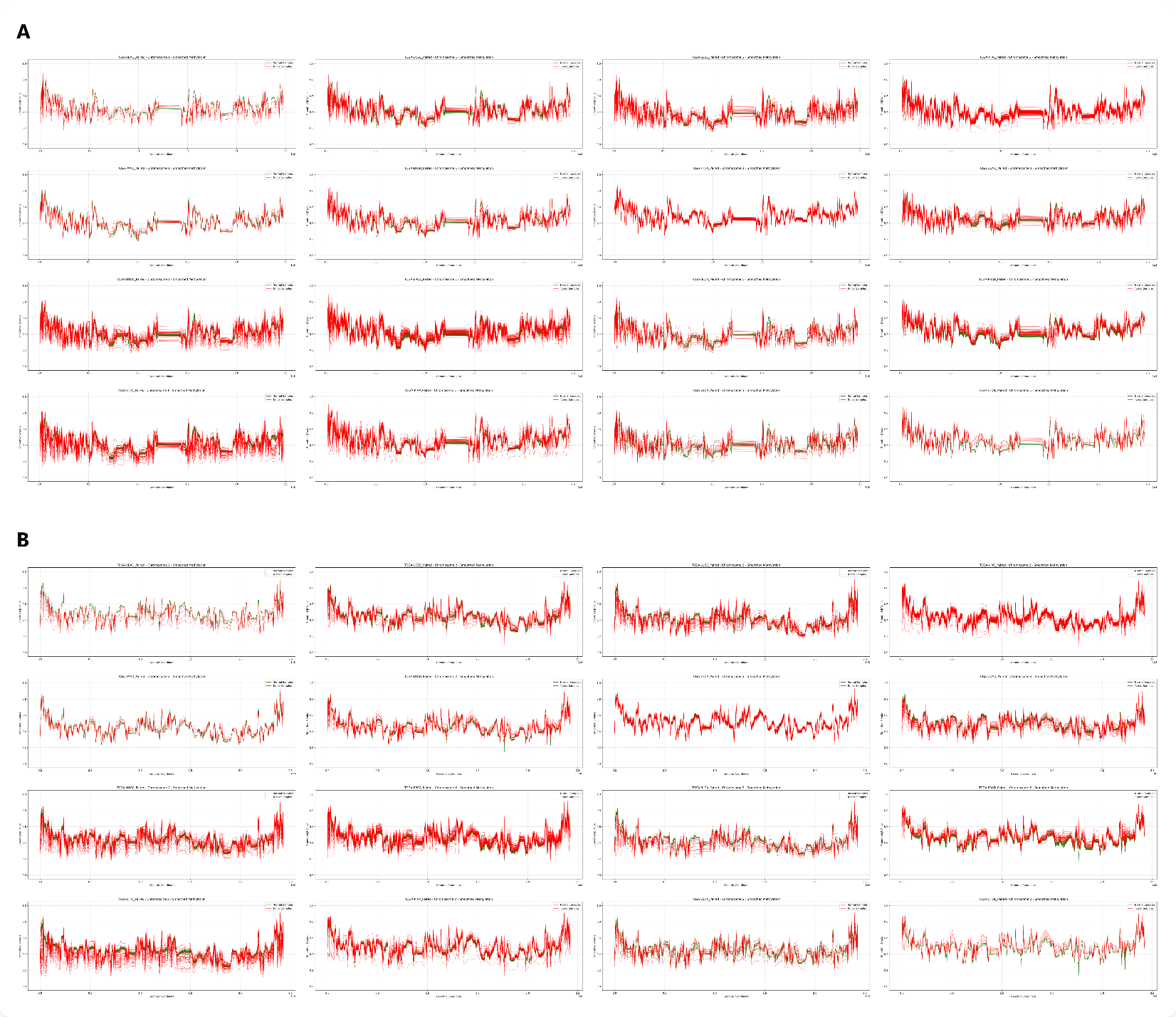
**Supplementary Figure 3. DNA methylation variability in paired tumor and normal samples from 16 types of cancer.** Smoothed chromosome-level methylation profiles across A) Chromosome 1, and B) Chromosome 2 for tumor and normal samples from 16 types of cancers, displayed for each tumor type separately. Similar results were obtained for Chr3-22 (not included in the manuscript).

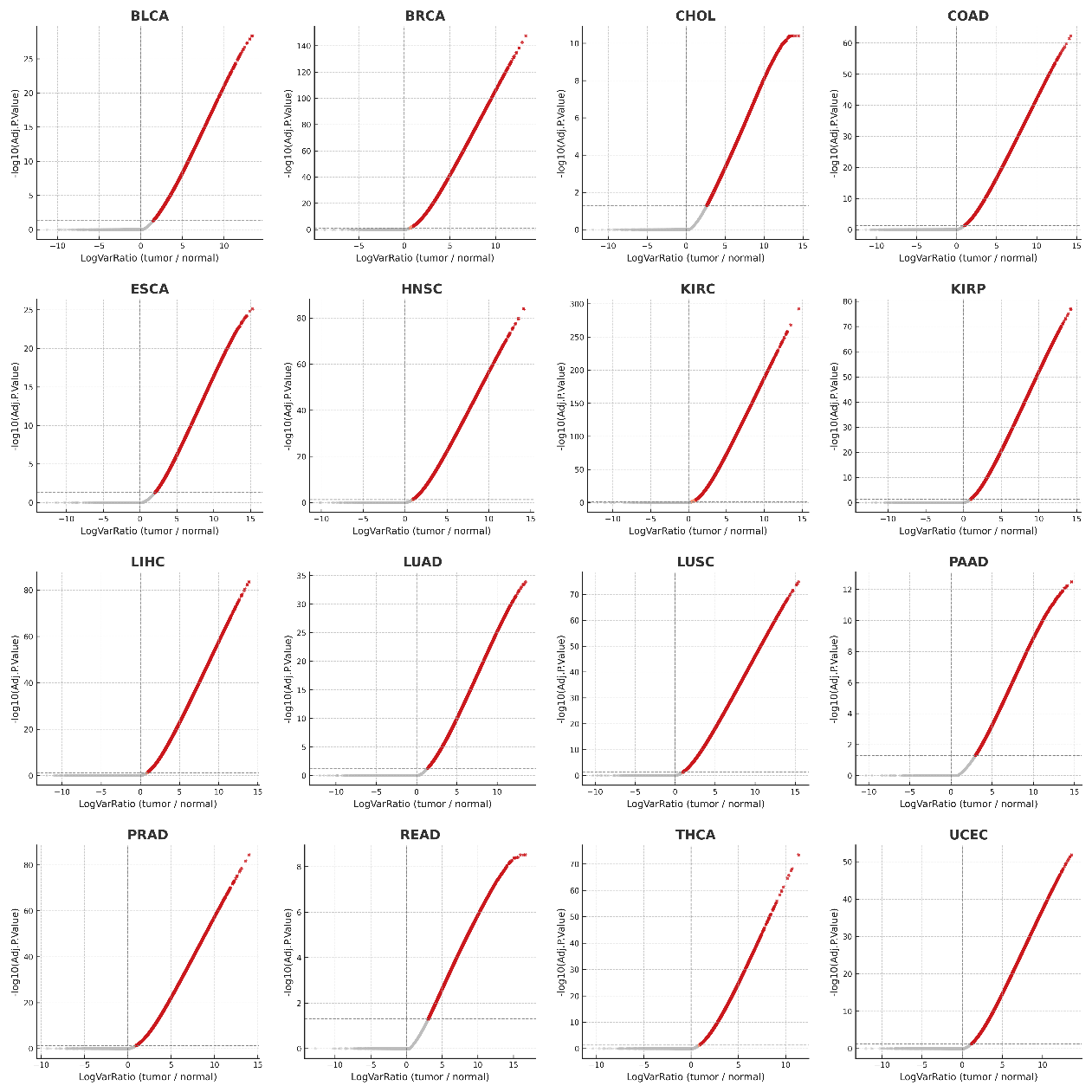

**Supplementary Figure 4. DNA methylation variability in paired tumor and normal samples from 16 types of cancer.** Volcano plot of log₂ variance ratios (tumor/normal) versus −log₁₀ Benjamini–Hochberg (FDR) adjusted P-values from differential variability testing.

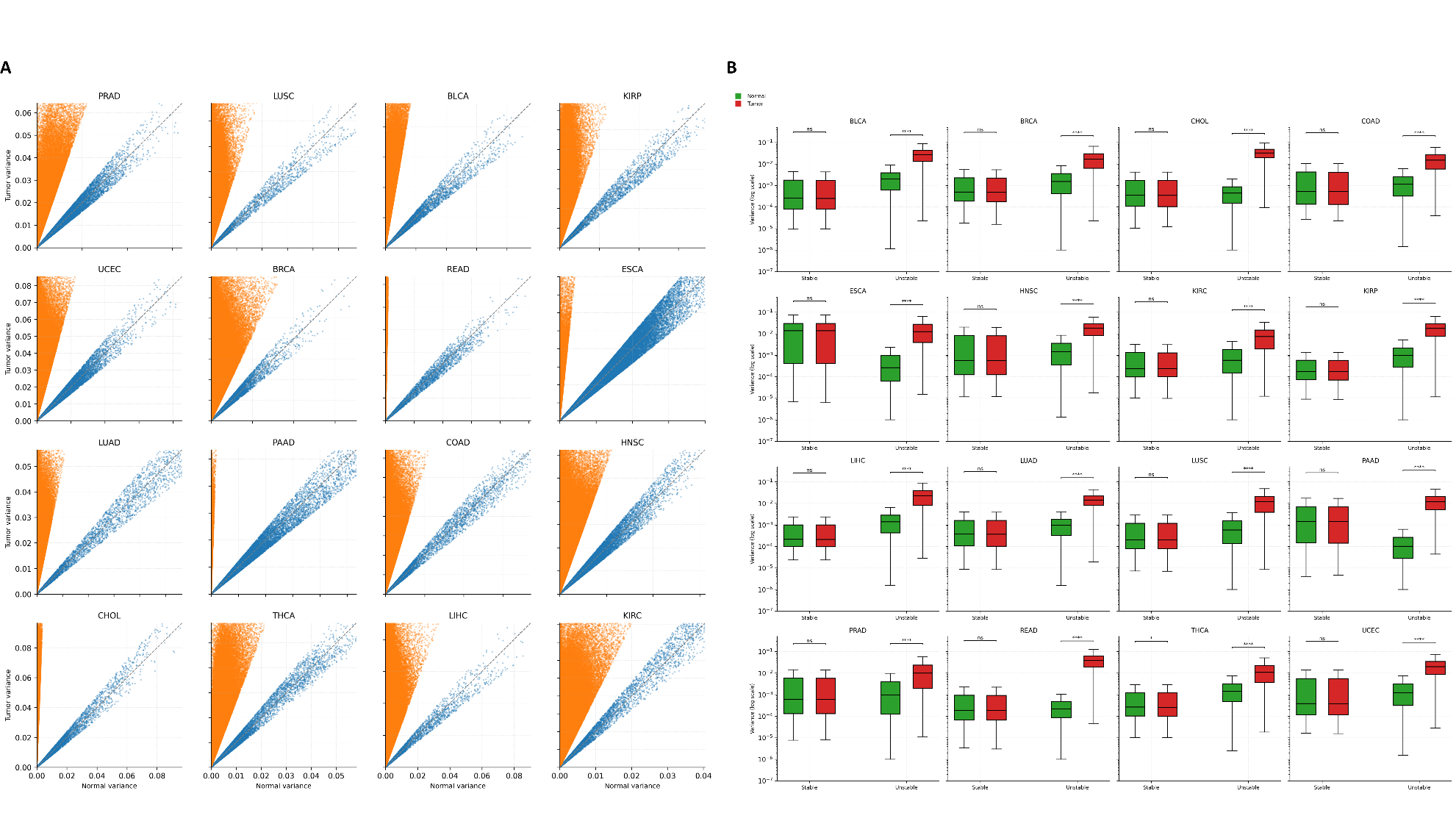
**Supplementary Figure 5. DNA methylation variability in paired tumor and normal samples from 16 types of cancer.** A) Scatter plot of tumor versus normal variance per CpG site, with points colored by stability class and the identity line shown for reference. B) Boxplots showing methylation variance for CpG sites classified as stable or unstable, displayed separately for tumor and normal samples on a logarithmic scale. Tumor–normal variance distributions within each class were compared using two-sided Mann–Whitney U tests on log₁₀-transformed variance values, with statistical significance.

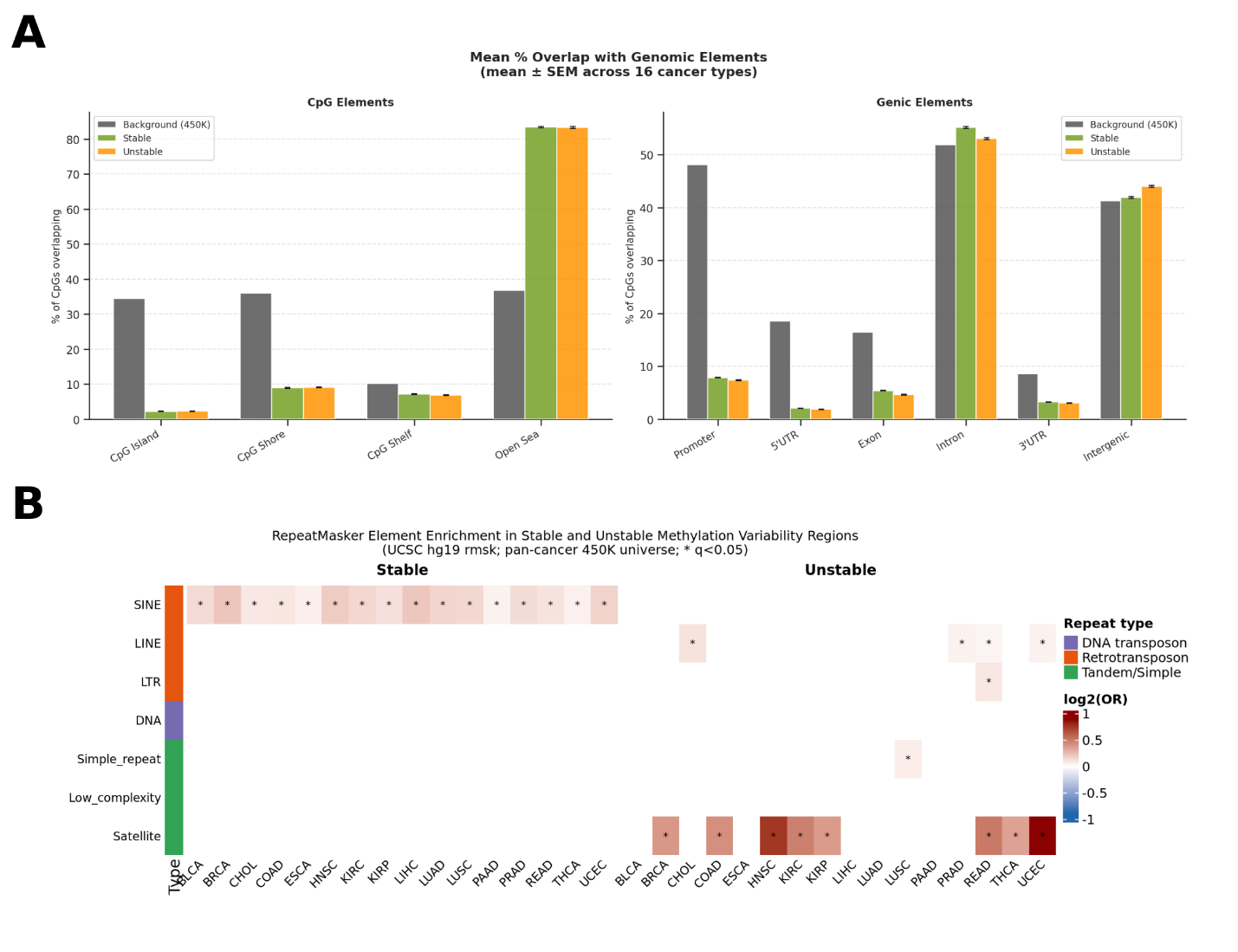

**Supplementary Figure 6. Genic, CpG, and repeat enrichment of stable and unstable DNA methylation regions across cancer types.** A) Bar plots summarizing enrichment of genic and CpG elements in stable and unstable regions across cancer types, B) Heatmap summarizing enrichment of repeat elements in stable and unstable regions across cancer types.

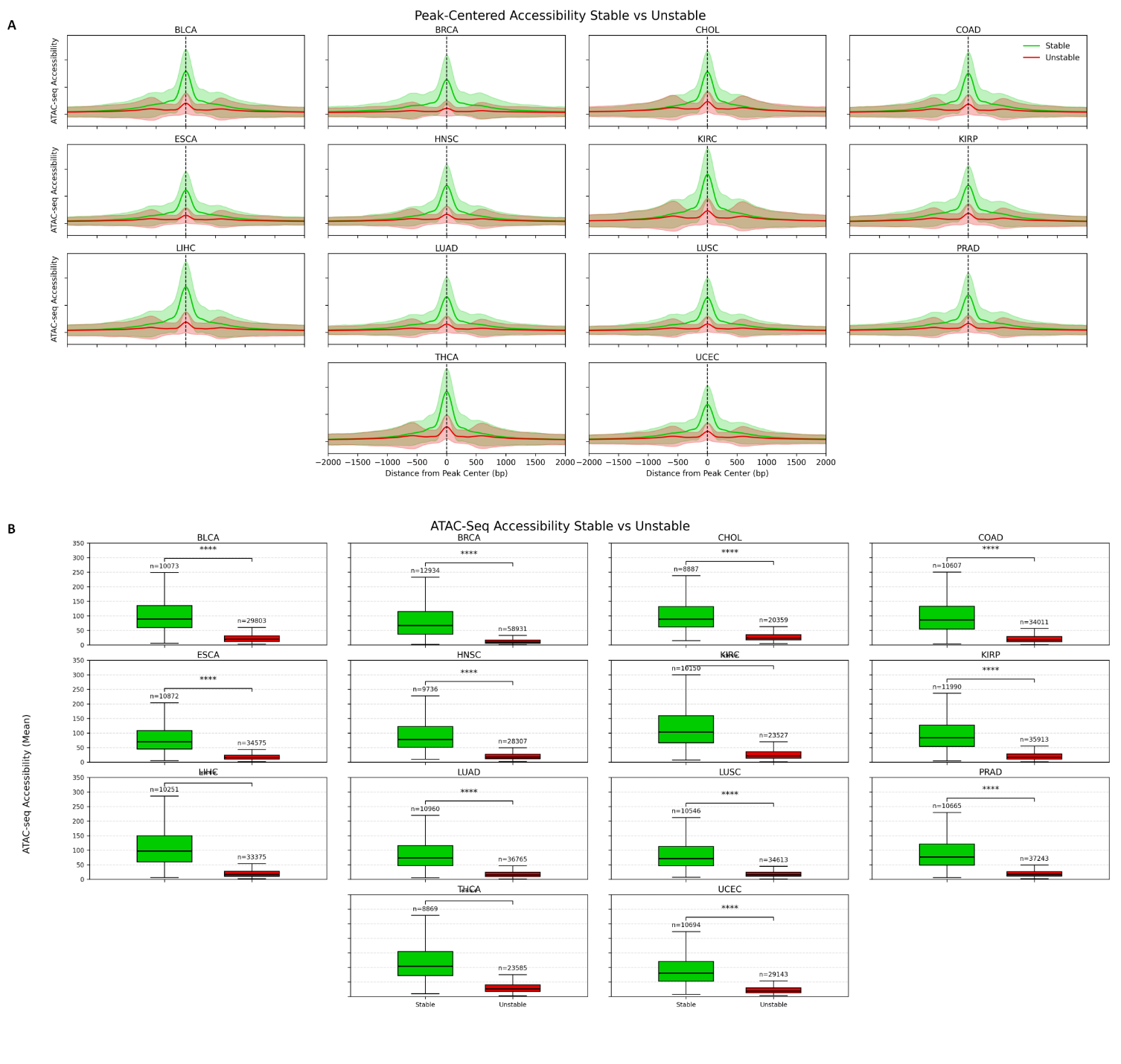
**Supplementary Figure 7. DNA methylation variability in tumor and normal samples from 16 types of cancer and their relationship with DNA accessibility.** A) Peak-centered ATAC-seq accessibility profiles for CpG regions classified as stable (green) or unstable (red) based on DNA methylation variability. Curves represent the mean normalized ATAC-seq signal centered on the region midpoint, with shaded areas indicating variability ±1 standard deviation (SD) across regions at each genomic position. The x-axis denotes distance from the peak center (±2 kb), and the y-axis denotes normalized chromatin accessibility, B) Boxplots summarizing mean ATAC-seq accessibility at stable and unstable regions for each cancer type. Each boxplot displays the distribution of region-level accessibility values, with sample sizes (number of regions) indicated. Statistical comparisons between stable and unstable regions were performed using a two-sided Mann–Whitney U test (Wilcoxon rank-sum test).

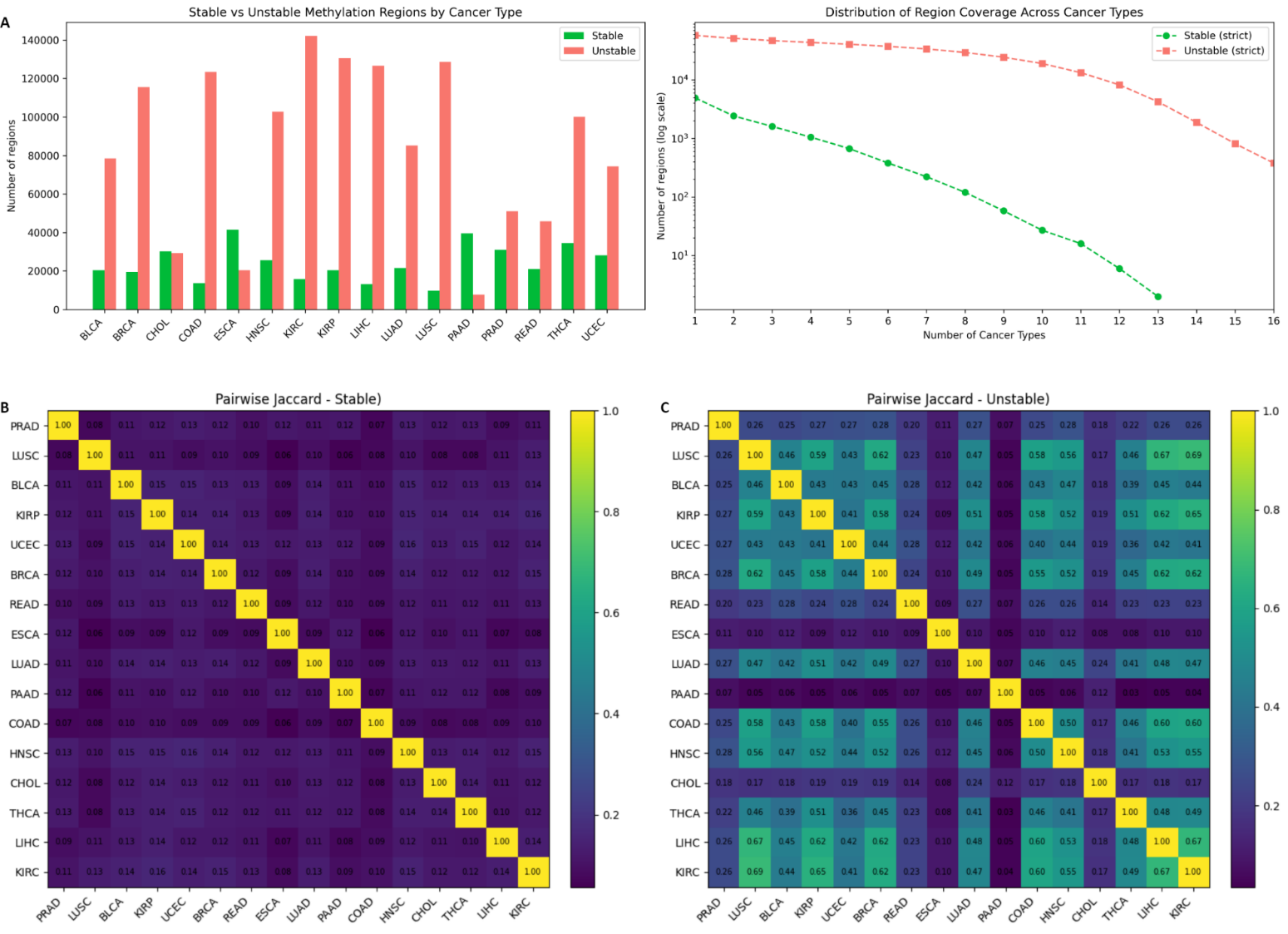

**Supplementary Figure 8. Definition of pan-stable and pan-unstable DNA methylation regions.** A) Bar plot showing the number of genomic regions classified as stable (green) or unstable (red) DNA methylation variability regions in each TCGA cancer type (left). Coverage curves summarizing the number of regions supported by increasing numbers of cancer types (k = 1–16) under a strict, conflict-free definition. The y-axis is shown on a logarithmic scale; green indicates pan-stable regions and red indicates pan-unstable regions (right). B) Heatmap of pairwise Jaccard similarity indices for stable regions across cancer types. C) Heatmap of pairwise Jaccard similarity indices for unstable regions across cancer types. Color scales indicate the magnitude of the Jaccard index, with diagonal elements fixed at 1.0.

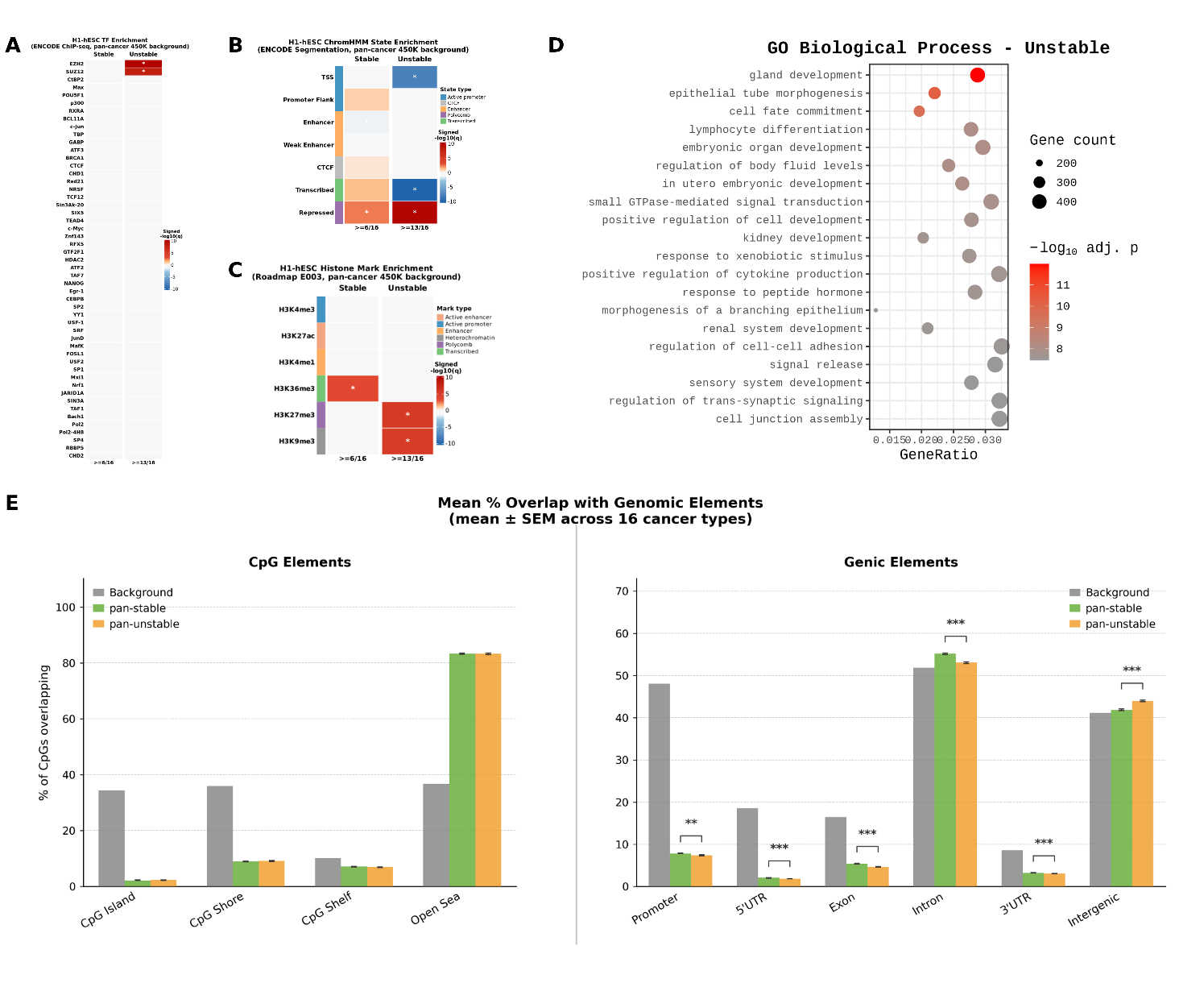

**Supplementary Figure 9. Regulatory and chromatin enrichment of pan-stable and pan-unstable DNA methylation regions.** A) Heatmap summarizing recurrent transcription factor (TF) enrichments for pan-stable and pan-unstable regions, B) Heatmap summarizing recurrent chromatin segment enrichments for pan-stable and pan-unstable regions, C) Heatmap summarizing recurrent histone modifications enrichments across cancer types for pan-stable and pan-unstable regions, D) Gene ontology analysis of enriched TF motifs and their target gene sets in pan-unstable (14/16) DNA methylation regions, E) Bar plot summarizing enrichment of pan-stable and pan-unstable regions (14/16) in genic and CpG elements.

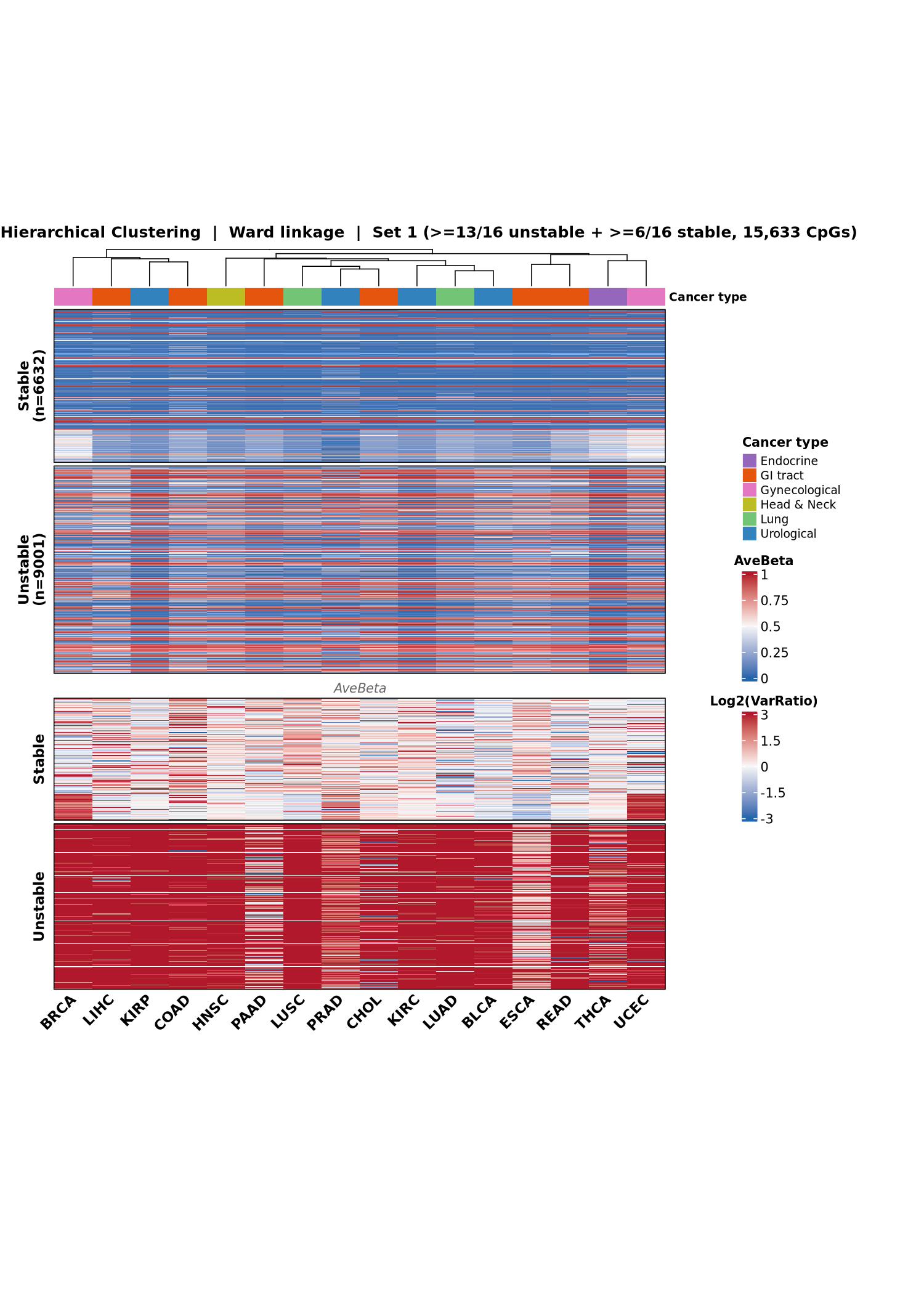
**Supplementary Figure 10. Cross-cancer clustering of pan-stable and pan-unstable DNA methylation sites.** Hierarchical clustering of DNA methylation across multiple cancer types using Ward linkage based on 15,633 CpG sites that were classified as either consistently stable (≥6/16 cancer types) or unstable (≥13/16 cancer types). Columns represent cancer types, and rows represent CpG loci. The upper heatmap shows average beta values (AveBeta) for each CpG across cancers, while the lower heatmap shows the log2-transformed variance ratio (log2(VarRatio)), highlighting differences in methylation variability between stable and unstable CpGs. CpGs are grouped into stable (n = 6,632) and unstable (n = 9,001) categories. The color scale in the top heatmap represents DNA methylation beta values (0–1), while the bottom heatmap shows log2-transformed variance ratios (−3 to 3), where red indicates higher variability and blue indicates lower variability. The annotation bar above the heatmap indicates the cancer system category.

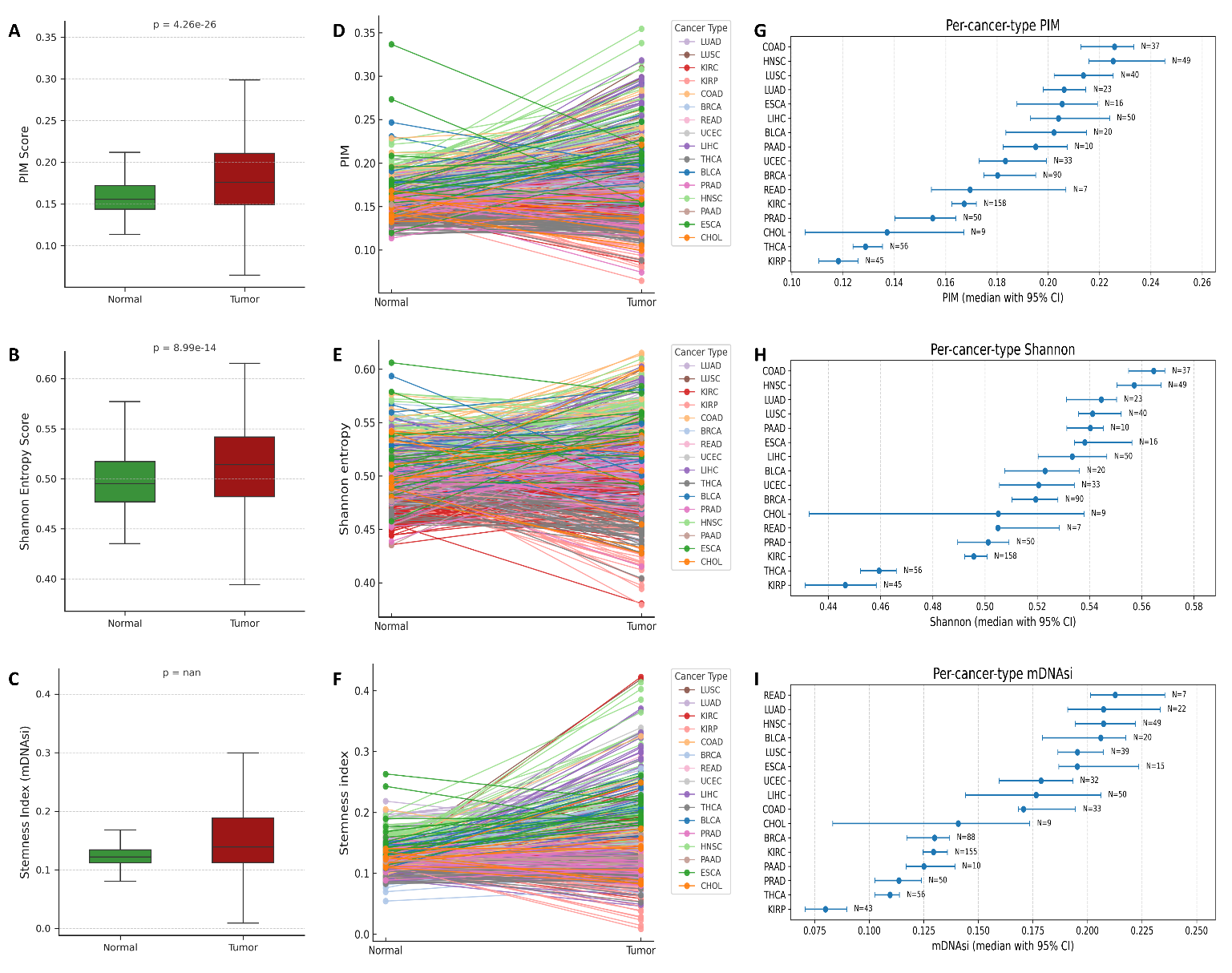

**Supplementary Figure 11. Genome-wide DNA methylation disorder across cancer types quantified by Proportion of Intermediately Methylated sites (PIM), Shannon entropy, and methylation-derived stemness (mDNAsi).** Boxplots comparing A) PIM, B) Shannon entropy, and C) mDNAsi for aggregated tumor (red) and matched normal (green) sample data. Statistical significance was calculated with the Wilcoxon paired test. Spaghetti plots of D) PIM, E) Shannon entropy, and F) Stemness index - mDNAsi matched tumor–normal pairs per cancer type, with each line representing an individual patient. Forest plot median values with 95% confidence intervals for G) PIM, H) Shannon entropy, and I) mDNAsi per cancer type.

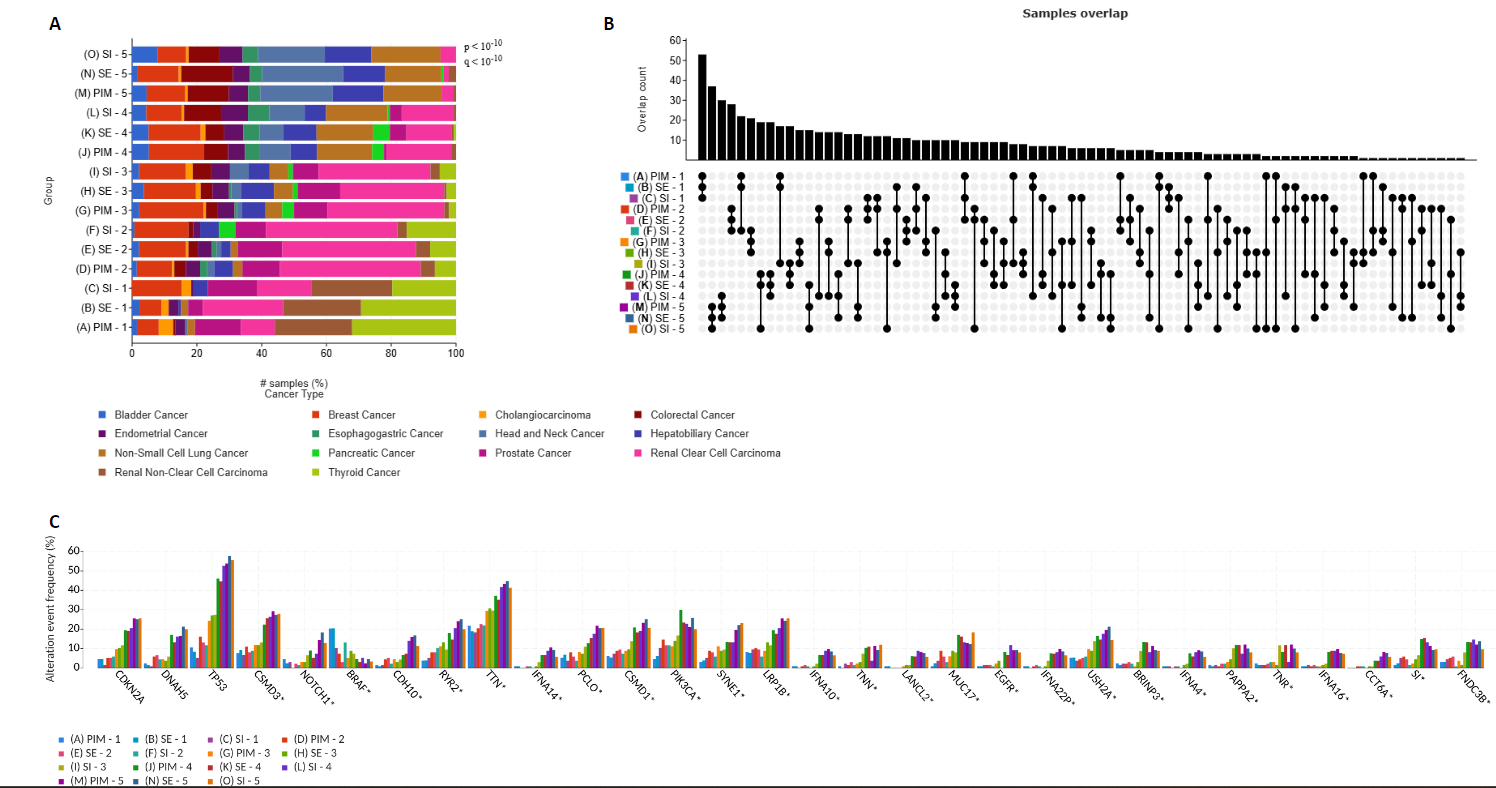
**Supplementary Figure 12. Co-stratification of tumors by DNA methylation disorder metrics and associated cancer-type and mutational profiles.** A) Stacked bar plots showing the cancer-type composition of samples within quintiles (1–5) of the three DNA methylation disorder metrics: PIM, Shannon entropy (SE), and methylation-derived stemness index - mDNAsi (SI). Bars represent the percentage of samples contributed by each cancer type within each quintile. B) UpSet plot depicting sample overlap across corresponding quintiles of PIM, SE, and SI, illustrating the extent of shared samples among disorder strata defined by different metrics. C) Bar plot showing the frequency of selected recurrent somatic gene alterations across methylation disorder quintiles for PIM, SE, and SI. Alteration frequencies are plotted per gene, stratified by quintile and disorder metric, enabling comparison of mutational patterns across increasing levels of DNA methylation disorder.

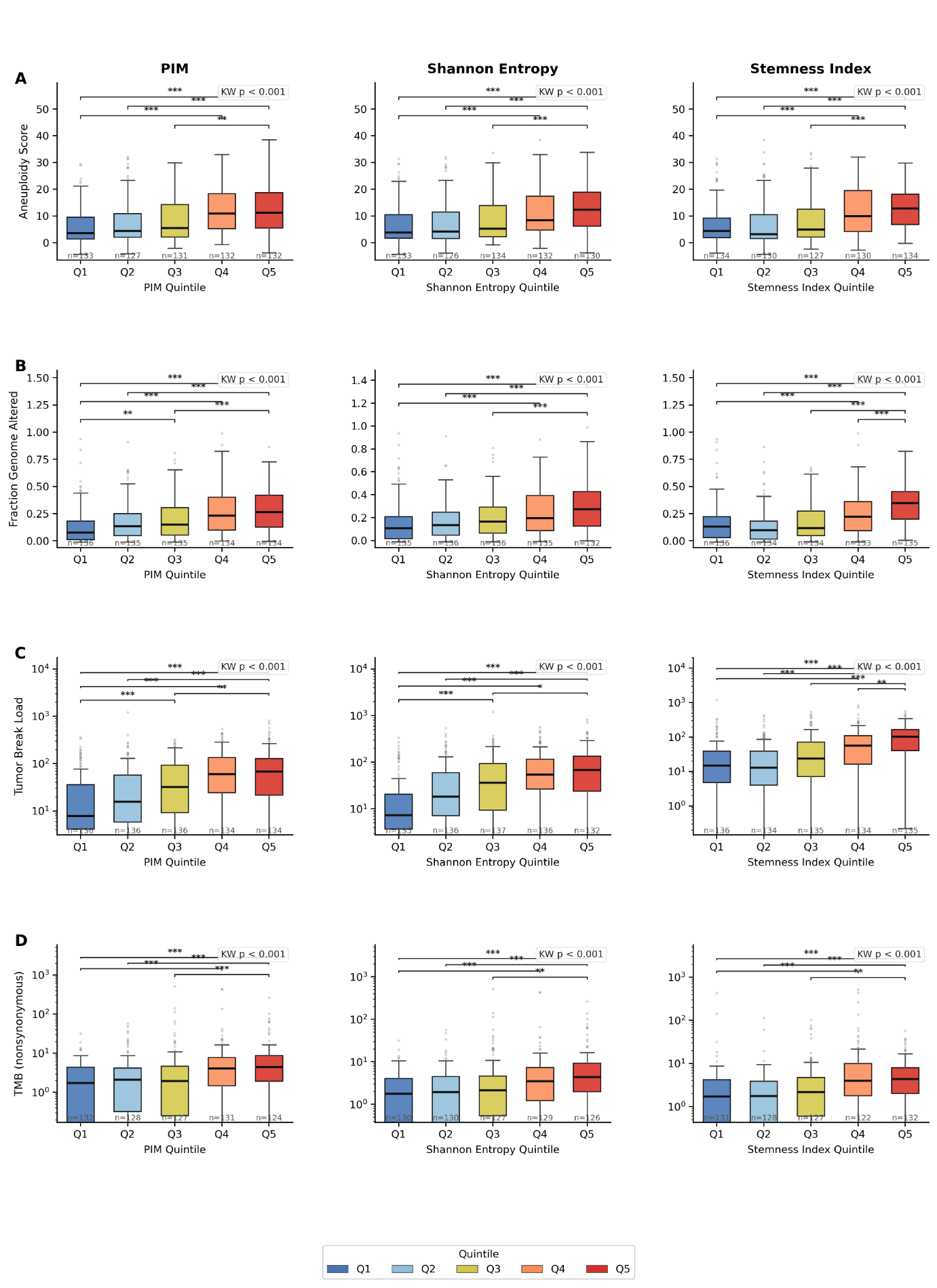

**Supplementary Figure 13. Association between DNA methylation disorder metrics and genomic instability features**. Boxplots of tumors stratified into quintiles (1–5) based on increasing levels of DNA methylation disorder as quantified by the Proportion of Intermediately Methylated sites (PIM), Shannon entropy, and methylation-derived stemness index. Boxplots show the distribution of genomic instability measures across quintiles for A) aneuploidy score, B) fraction of genome altered (FGA), C) tumor break load, and D) tumor mutational burden (TMB), adjusted for tumor purity. Boxes represent the interquartile range with median values indicated; whiskers denote 1.5× IQR, and points represent outliers. Brackets indicate comparisons across increasing quintiles within each metric, with statistical significance calculated using the Mann–Whitney U test.

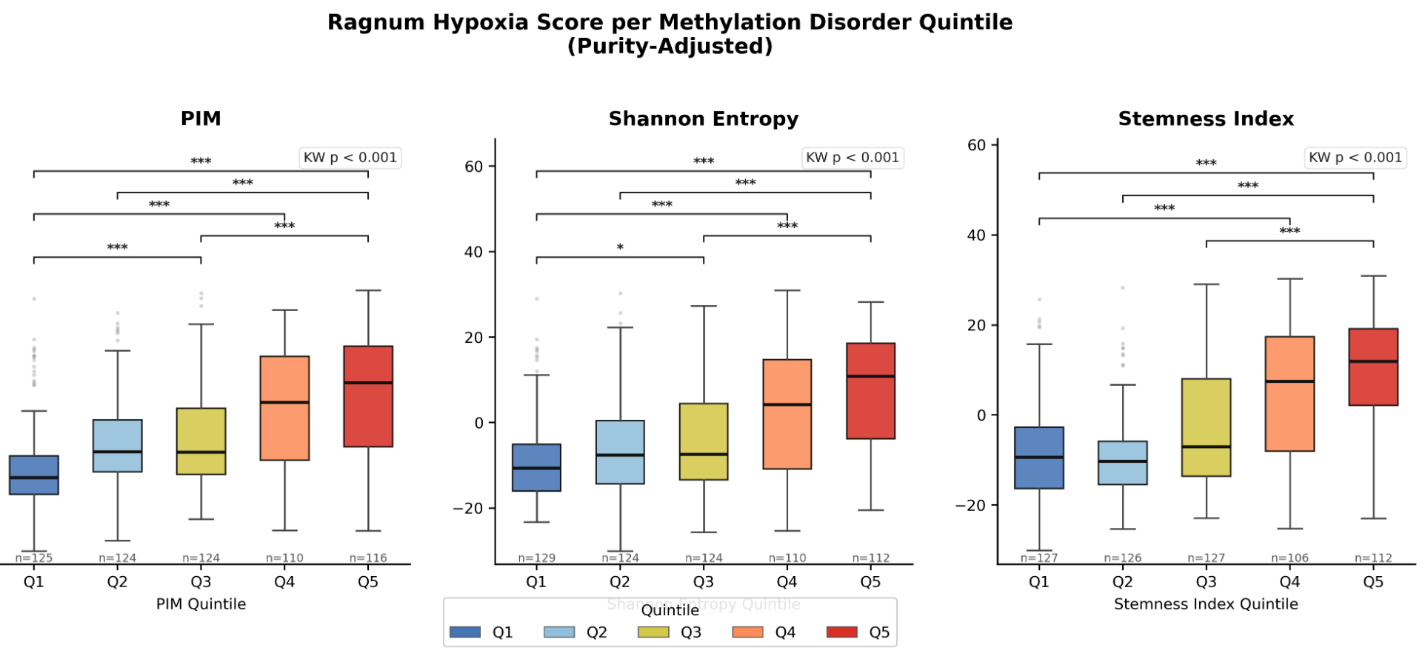

**Supplementary Figure 14. Association between DNA methylation disorder metrics and Ragnum hypoxia score**. Boxplots of tumors stratified into quintiles (1–5) based on increasing levels of DNA methylation disorder as quantified by PIM score (left), Shannon entropy (middle), and stemness index mDNAsi (right), adjusted for tumor purity. Boxes represent the interquartile range with median values indicated; whiskers denote 1.5× IQR, and points represent outliers. Brackets indicate comparisons across increasing quintiles within each metric, with statistical significance calculated using the Mann–Whitney U test.

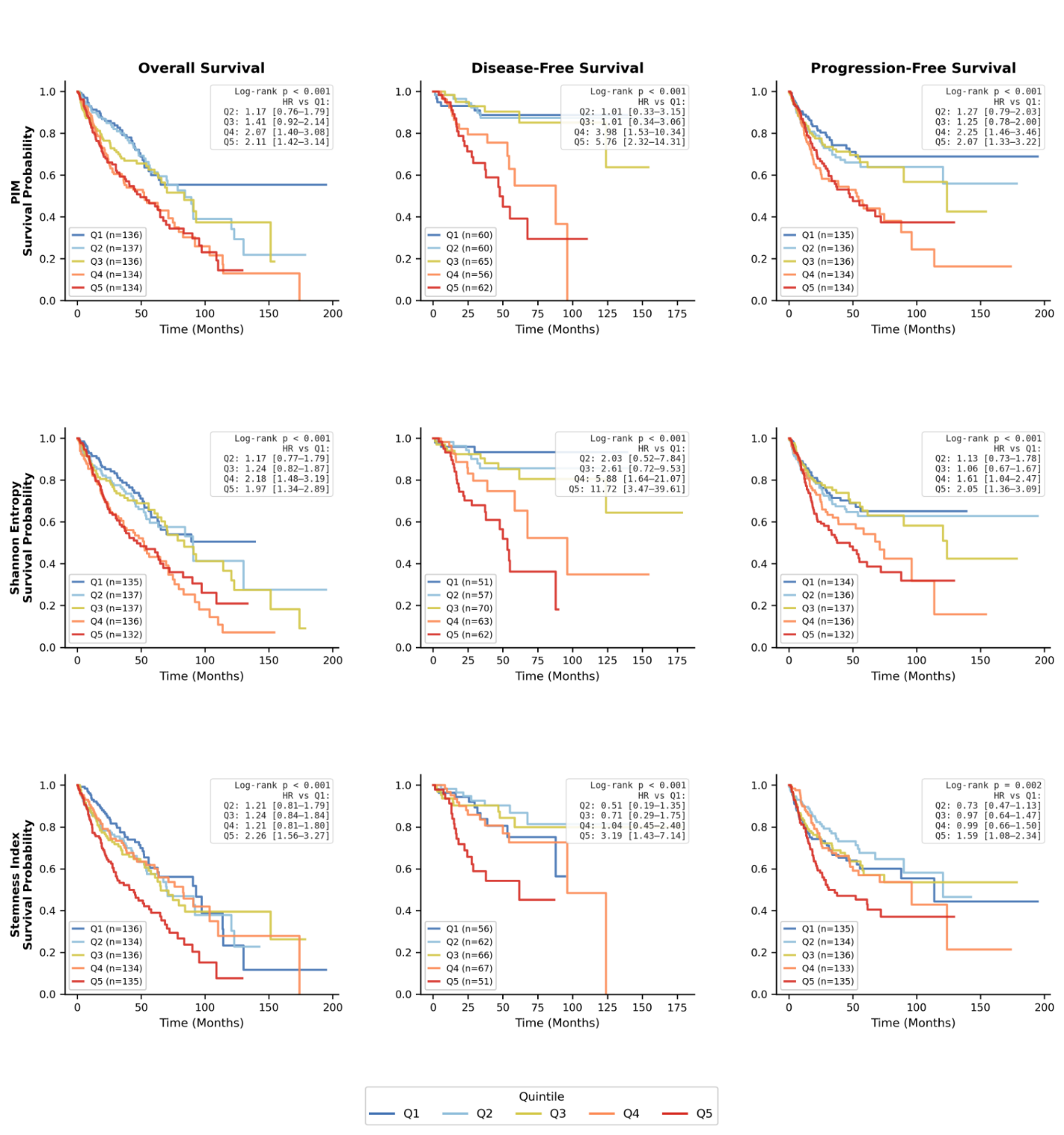

**Supplementary Figure 15. Survival analyses stratified by DNA methylation disorder metrics**. Kaplan–Meier survival curves are shown for overall survival (left), disease-free survival (middle), and progression-free survival (right), adjusted for tumor purity, with patients stratified into quintiles (1–5) according to increasing levels of DNA methylation disorder as quantified by the proportion of intermediately methylated sites (PIM), Shannon entropy, and the methylation-derived stemness index mDNAsi. Curves are color-coded by quintile. Log-rank test P values for each comparison are indicated within panels. Hazard ratios with 95% confidence intervals for each quintile relative to the lowest-disorder group are reported in the legends of the respective panels.

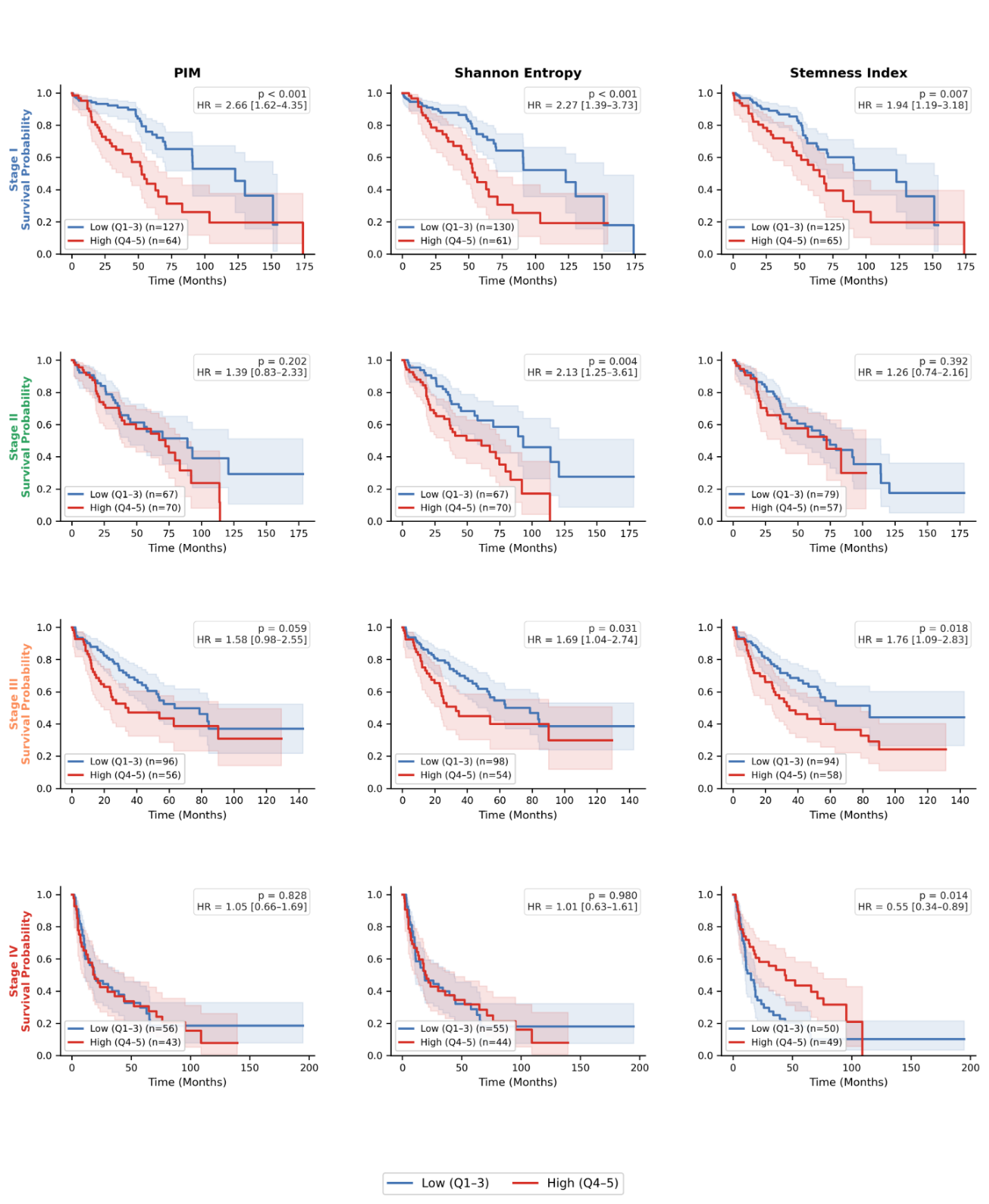

**Supplementary Figure 16. Overall survival analyses stratified by DNA methylation disorder metrics and staging.** Kaplan–Meier survival curves are shown PIM score (left), Shannon entropy (middle), and stemness index mDNAsi (right), adjusted for tumor purity, with patients stratified into low disorder metric (quintiles 1–3), and high disorder metric (quintiles 4-5). Curves are color-coded by the status of the disorder metric. Log-rank test P values for each comparison are indicated within panels. Hazard ratios with 95% confidence intervals for high relative to the low-disorder group are reported in the legends of the respective panels.
